# High precision automated detection of labeled nuclei in Gigapixel resolution image data of Mouse Brain

**DOI:** 10.1101/252247

**Authors:** Sukhendu Das, Jaikishan Jayakumar, Samik Banerjee, Janani Ramaswamy, Venu Vangala, Keerthi Ram, Partha Mitra

**Affiliations:** Department of Computer Science & Engineering, Indian Institute of Technology Madras, Chennai, India; Center for Computational Brain Research, Indian Institute of Technology Madras, Chennai, India; Healthcare Technology Innovation Centre, Indian Institute of Technology Madras, Chennai, India; Cold Spring Harbor Laboratory, NY, USA

**Keywords:** GFP, cell detection, convex arc, mouse brain, distance transformation, ridge

## Abstract

There is a need in modern neuroscience for accurate and automated image processing techniques for analyzing the large volume of neuroanatomical imaging data. Even at light microscopic levels, imaging mouse brains produces individual data volumes in the TerraByte range. A fundamental task involves the detection and quantification of objects of a given type, e.g. neuronal nuclei or somata, in brain scan dataset. Traditionally this quantification has been performed by human visual inspection with high accuracy, that is not scalable. When modern automated CNN and SVM-based methods are used to solve this classification problem, they achieve accuracy levels that range between 85 – 92%. However, higher rates of precision and recall that are close to that of human performance are necessary. In this paper, we describe an unsupervised, iterative algorithm, which provides a high performance for a specific problem of detecting Green Fluorescent Protein labeled nuclei in 2D scans of mouse brains. The algorithm judiciously combines classical computer vision techniques and is focused on the complex problem of decomposing strong overlapped objects of interest. Our proposed technique uses feature detection methods on ridge lines over distance transformation of the image and an arc based iterative spatial-filling method to solve the problem. We demonstrate our results on mouse brain dataset of Gigabyte resolution and compare it with manual annotation of the brains. Our results show that an aptly designed CV algorithm with classical feature extractors when tailored to this problem of interest achieves near-ideal human-like performance. Quantitative comparative analysis, using manually annotated ground truth, reveals that our approach performs better on mouse brain scans than general purpose machine learning (including deep CNN) methods.

## I. Introduction

Understanding the cytoarchitecture of the brain is an important goal in the field of neuroscience [26], [52]. In particular, the identification of locations and quantification of neurons across the brain is a vital task. To achieve this goal, modern imaging methods have allowed users to produce images at very high resolution for further analysis. Entire mouse brains can now be imaged using light microscopy at sub-micron resolution from entire mouse brains (~ 1*cm*^3^ volume), with individual data volumes in the terrabyte range. Optical microscopic techniques applied at such high resolutions, help to study the brain at a single neuron resolution [37], [43] but the large data volumes make it a significant computational and data analytics challenge.

In addition to the advancements in imaging techniques, advanced methods such as the expression of the Green Fluorescent Protein (GFP) [12] have ensured that specific types of neurons can be tagged and studied in great detail [55]. A successful implementation of GFP tagging in the identification and quantification of interneurons is described in [55]. Manually studying the distribution of these GFP expressing neurons across the entire brain is often time consuming and is susceptible to human errors over large datasets. To overcome this, an automated process of detection and quantification of neurons [5], [28] must be designed based on well grounded image processing (or computer vision) algorithms. However, the level of sophistication in the analysis techniques has not yet caught up with the significant improvements in microscopic imaging methods. In this work, we have sought to develop an automated method of detection for GFP tagged interneurons across mouse brain datasets that is made up of high resolution image scans of brain sections. Thus, the aim of our work, proposed in this paper, is to detect and count GFP tagged specific cell types from entire mouse brains.

Automated algorithms are often resource consuming and have been deployed in studying the brains of lower species such as Candida elegans [56]. Computer vision algorithms have also played a vital role in the segmentation of large-scale electron microscopy data. Kaynig *et al*. [27] have proposed a pipeline that provides state-of-the-art reconstruction performance while scaling to data sets in the GB-TB range. Such high performance was achieved by training a random forest classifier on interactive sparse user annotations. This classifier output was then combined with an anisotropic smoothing prior in a Conditional Random Field (CRF) framework to generate multiple segmentation hypotheses per image. These segmentations were then combined into geometrically consistent 3D objects by segmentation fusion. A more recent work in automatic neural reconstruction from petavoxel of Electron Microscopy data [53] suggested a dense Automatic Neural Annotation framework (RhoANA) to automatically align, segment and reconstruct a 1*mm*^3^ brain tissue (~ 2 peta-pixels), using a web-based tool to manually proofread the output, and ensure reconstruction correctness. The pipeline performs membrane classification and 2D segmentation using state-of-the-art deep learning (DL) techniques in order to generate membrane probability maps.

Deep learning methods have been used in the detection of cells in cancer biology. One method for mitosis detection [4] is based on deep learning which performs data aggregation using convolutional neural networks (CNN) via additional crowdsourcing layer (AggNet). Mualla *et al*. [39] propose a method for automatic cell detection on polypropylene substrate, suitable for microbeam irradiation. A Harris corner detector was employed to detect apparent cellular features. These feature-corners were grouped based on a dual-clustering technique according to the density of their distribution across the image. Weighted centroids were then extracted from the clusters of corners and constituted the targets for irradiation. Quelhas *et al*. [46] introduce a new approach for cell detection and shape estimation to detect overall convex shapes in multivariate images based on a sliding band filter (SBF). Furthermore, the parameters involved are intuitive as they are directly related to the expected cell size. Using the SBF filter, cell nuclei, the cytoplasm location and shapes were detected. In the remainder of the paper, we will refer to the terms cells and GFP nuclei interchangeably (as our target objects for detection/segmentation).

**Fig. 1:**
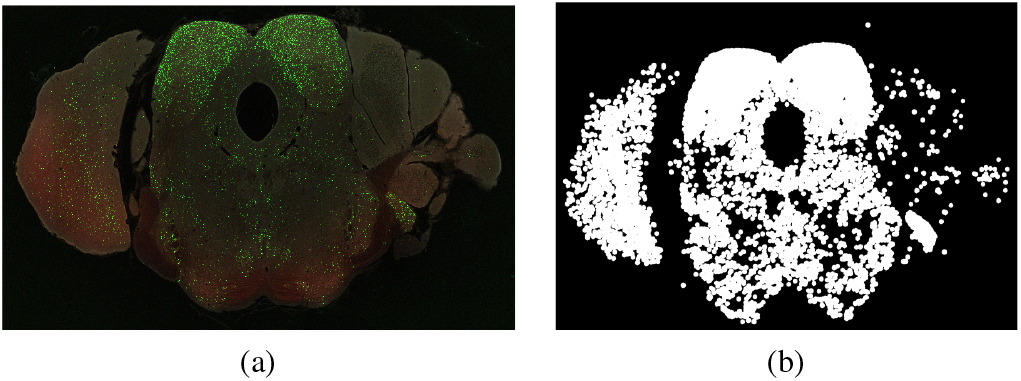
(a) Example of a high resolution image of an individual brain section highlighting the distribution of GFP expressing interneurons. This brain section image along with ~ 240 separate images constitutes one complete brain dataset. (b) shows the extracted Green channel data from (a).

An example of a high resolution image (2D) of a brain section is given in Figure 1. This brain section along with 240 brain sections form a complete brain imaged dataset of one particular mouse brain (*Mouse Brain Hua-167* [1]). The dataset consists of images with a 18*K* × 24*K* resolution at 0.46 micron per pixel scale. The given example shows the distribution of the GFP expressing nuclei with higher distribution densities clearly visible in the colliculi. Figure 1b shows the extracted image of the green channel alone for the same brain section. Figure 2a shows a zoomed in example of a small cropped region from the example section (figure 1). Centers of the GFP nuclei were marked manually by human annotators for the entire brain and this ground truth has been shown in Figure 2b. Results of the object-detection methods based on Faster R-CNN [50] and SVM [16], [40] have also been shown. As demonstrated in figures 2(c) and 2(d), although these methods yield fairly accurate results, however produce a few false alarms (detections) and misses, especially when detecting strong overlapping objects (Section V discusses performance using quantitative measures). As a scientific and diagnostic tool, there is a need for a higher level of accuracy and higher rates of precision and recall. The major reason for the moderate accuracy given by these supervised methods is that, they do not capture the various non-unique patterns of spatial layouts and structures that are formed by clusters of neurons, specifically with strong overlap. Machine learning algorithms [7], [20] have been designed to produce high accuracy invariant to pose, scale (size), background and texture of the object. However, they are not immune to conditions where large amount of overlap (dense crowd, large clutter/pile of objects, say) occur and fail to inadequately capture the required objects of interest due to occlusion and camouflage. Deep-CNNs have yielded rich dividends in example cases, ranging from IMAGNET [19], PASCAL [21] image datasets, to HMDB [29], Hollywood-II [34], MS-COCO [30] video datasets, where considerable inter-class variability exists in the data, unlike in our dataset. Moreover, none of these publicly available datasets contain objects with large amount of clutter and strong overlap, which is the case in some scans of our brain dataset.

**Fig. 2:**
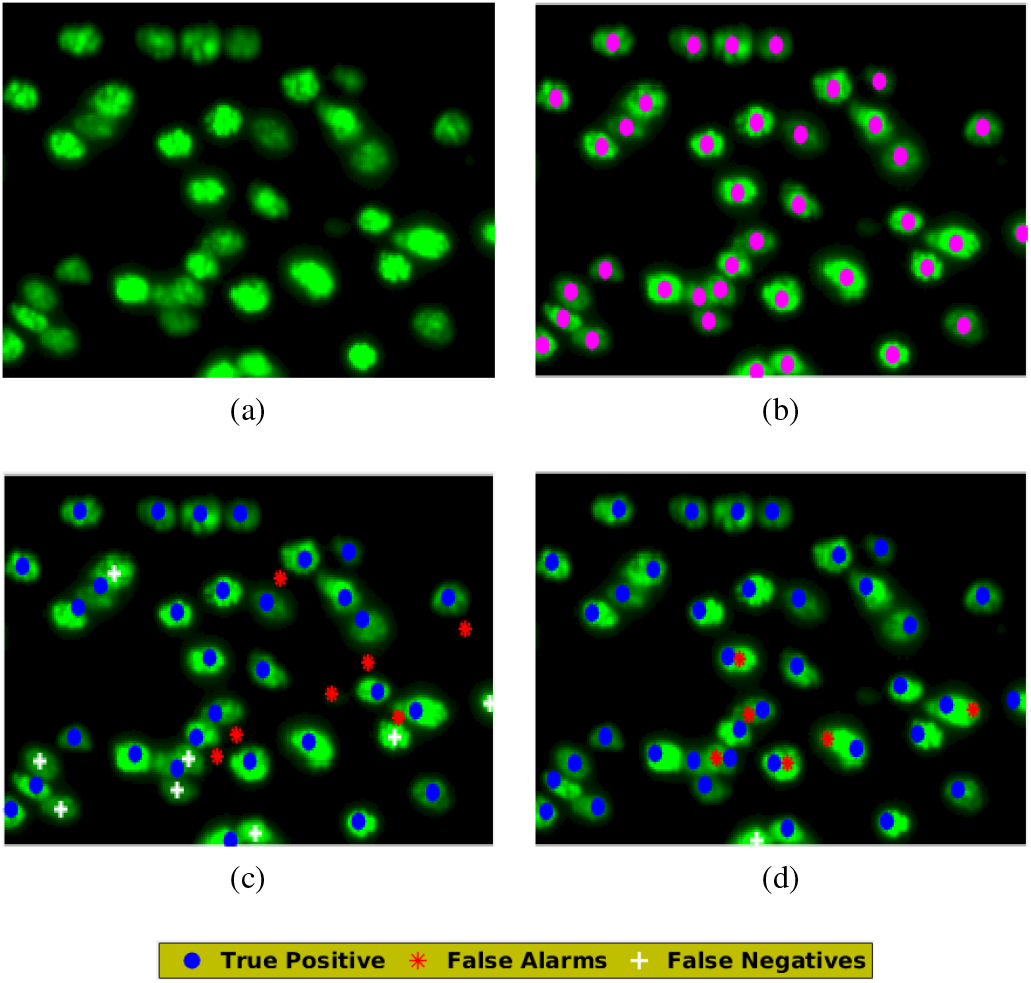
Applications of Support Vector Machines (SVM) and Faster R-CNN for the detection of GFP tagged nuclei on a mouse brain section; For clarity purposes, (a) cropped sample of the high resolution image is shown from brain section in figure 1(a); Manually annotated ground truth of cell centers for the sample shown in (a) (marked in magenta dots); Results of cell center detection using: (c) SVM [16] (Gaussian kernel) & (d) Faster R-CNN [50] on (a). Figure best viewed in color. Compared with the ground truth, the true positives are shown in blue, the false alarms in red and the false negatives in white crosses (consistently followed henceforth).

Detection of cell centers in cases of strong overlap of objects of interest is a difficult problem, as the overlapping objects result in complex 2D structures that do not exhibit a unique appearance pattern of shape, size and overlap formation (layout). This results in loss of accuracy in the training of any shallow or deep learning (DL) methods. However, human observations can intuitively sort these overlapping objects and it becomes necessary for any automated algorithm to achieve high accuracy close to that of human annotators but over a large scale (≃ 400 GB size images) with lesser computational time (few minutes on a single high-end CPU machine). With this aim, we have proposed an improved and efficient algorithm for the unsupervised classification of GFP tagged GABAergic interneurons across the mouse brain [8]. Our proposed method is designed suitably as a congruence of image analysis algorithms, such as Distance Transformation (DT) and edge detection to quantify the boundaries of the detected cells, followed by the extraction of features along ridge lines for detecting the candidate centers of neurons. The algorithm is further enhanced by an iterative approach of region filling by circular blobs of adaptive size, using convex arcs on the edge maps for detecting overlapped cell areas. Using this method, we have obtained a high precision of 0.972 and a recall of 0.961, over one complete brain section image set and a significant part of another (~ 240 scans in first set and ~ 158 in the other). A comparative performance analysis shows that our proposed algorithm surpasses two existing deep learning CNN models [50], as well as two popular shallow learning object detection algorithms. Shallow and deep models, used in this study, are not tailor-made for a structure decomposition problem which is an essential component necessary for detecting overlapping objects (GFP tagged nuclei in our case).

## II. Overview of the Proposed Method

Individual sections of the mouse brain are obtained as gigapixel resolution (RGB 18*K* × 24*K* pixels at 12-bits per channel) images, from the mouse brain of the wild type mouse (C57BL6), with green fluorescent protein (GFP) tagging of GABAergic neurons (PV+) [1]. This data is part of the mouse brain architecture project [1] at the Cold Spring Harbor Laboratory (CSHL), which aims to quantify the connections of different parts of the brain using tracer injections and cell tagging techniques at a mesoscopic scale [8]. The CRE cell lines that tag interneurons were developed by Josh Huang’s Laboratory and have been described in [55]. The histological and imaging preparation of the brain has been previously described in [45]. Manual annotation of one complete set of a brain and a selected portion of another set was performed by researchers from the Indian Institute of Technology (IIT), Madras, India, making the comparison and evaluation of algorithms feasible on the scan data. Details of the manual annotation is described later in Section V.

Figure 3 shows two examples of zoomed in sections with GFP tagged neurons with different cell distributions (as density of occurrence) - minimal overlap and strong overlap. Objects of interest (nuclei), which are visible in the green spectrum, in the strong overlapping regions appear as a dense 2D packed structure with no well formed/defined shape and size, where more than 3 cells overlap (figure 3(b)). In addition, individual nuclei also appear with varying sizes within the regions of interest. This arbitrary spatial arrangement of cells with significant overlap makes it a challenging task to detect each one of them accurately. The unsupervised computer vision (CV) approach of cell detection proposed in this paper has been designed and formulated for precisely dealing with this scenario of detecting individual cells within a packed set of strongly overlapping nuclei. Geometric constraints of the cell layout have been used to decompose a structure into overlapping circular units.

**Fig. 3:**
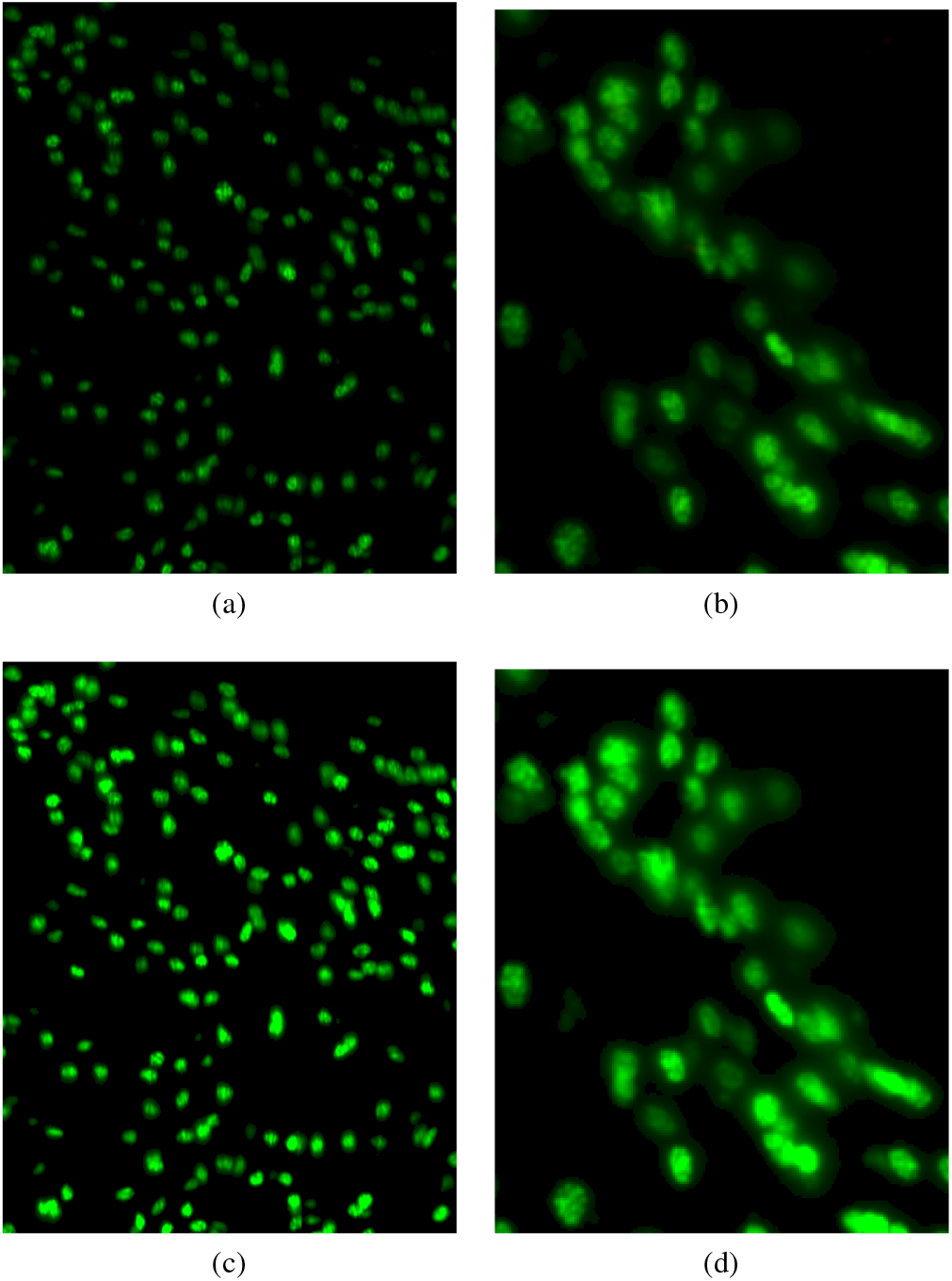
Examples of distributions of GFP tagged interneurons in a mouse brain, for (a) Minimal overlap, (b) Strong overlap; (c), (d) Extracted salient foreground region (FGR) obtained by enhancing the contrast of (a) and (b) respectively (best viewed in color). Other non-green channels are suppressed. See Supplementary section S.1 for details and illustration of the process of contrast enhancement.

### A. Basis and Justification of the algorithm designed

For our problem scenario, the task involves detecting centers of the GFP tagged nuclei. Our solution is achieved by using Distance Transformation and edge detection followed by Ridge estimation as default processes of *Stage I* of our pipeline. This part of the designed process accurately detects cells in cases of sparse (isolated) distribution or minimal overlap of objects of interest. In the complex case of strong overlap of multiple cells (as seen in figure 3(b)), an additional process (*Stage II*) is required for accurate identification of the nuclei. This solution is where our algorithm considerably outperforms the state of the art machine learning models (like Faster R-CNN [50], SVM [16]). The process we have designed uses geometric constraints with the foreground area filling to decompose a 2-D area of overlapping cells into partly occluded cell bodies. The complexity in the data lies in the degree of overlap of objects within the image and is dependent on the type of object, the microscopic and sample preparation techniques. The deep learning models, although well suited for multiple object detection are not well suited to the overlap-decomposition problem which we have implicitly addressed in our pipeline. The simplicity of method when combined with close to human performance, makes it advantageous to use over other methods including CNNs for solving this problem.

**Fig. 4:**
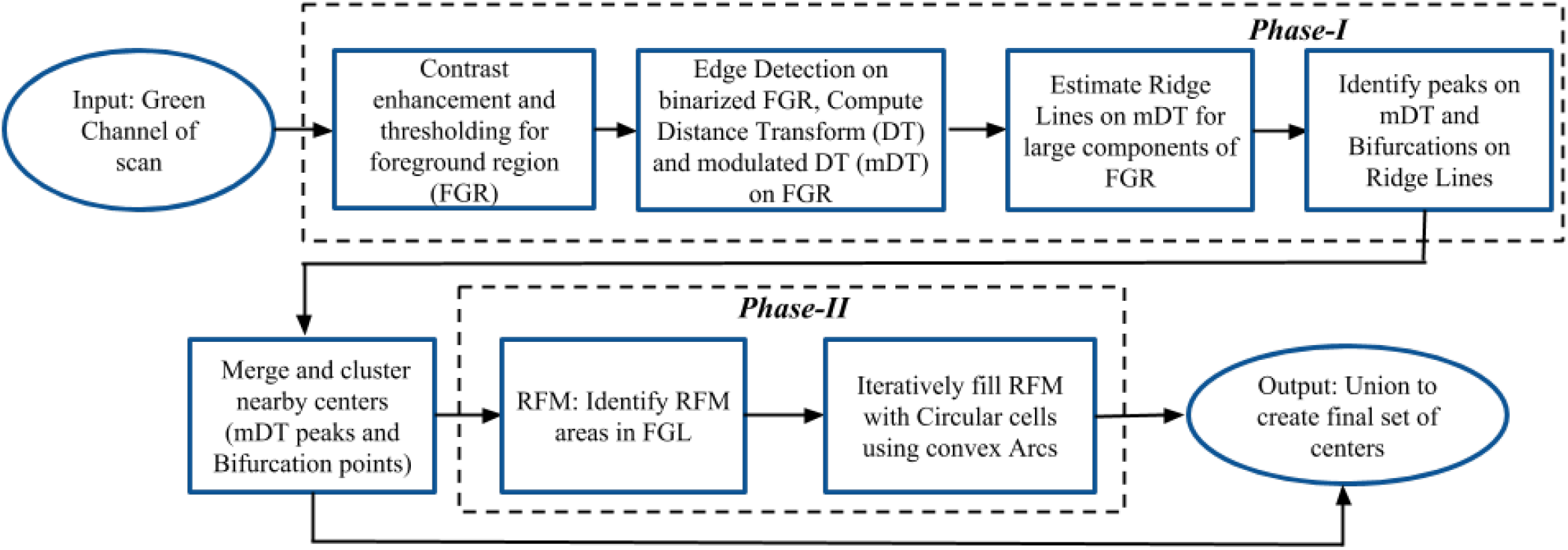
Flowchart detailing the algorithmic flow of the proposed method for the detection of centers of GFP tagged nuclei in mouse brains. *Phase I*: Method based on Distance Transform and ridge based features; *Phase II*: the arc based iterative method of area-filling. A more detailed outline of *Phase II* is given in algorithm 1 (and figure S.2).

### B. Brief Description of the proposed algorithm

As the GFP tagged neurons have relatively higher irradiance in green channel compared to the background neuropil, we have enhanced the 12-bit green channel using contrast enhancement as a pre-processing step to produce the salient foreground region (FGR) as shown in figures 3(c) and 3(d).

It was observed that modern machine learning (ML) and deep learning (DL) based supervised object detection algorithms fail to reach the desired level of high accuracy in the case of strong overlapping (figure 3(b)) arrangement of cells (results illustrated in supplementary section S.6). This task is analogous to the problem of packing circles of varying (but within a limited range) sizes within an arbitrary polygonal shape. Simpler versions of this problem have been addressed in the fields of Computational Geometry and Operational Research [32], [48], [25], [35], [15], [11], [58], [33], [17]. This is an NP-hard problem [14] and no prior work exists to give a direct solution for such a task, specifically in cases of overlap of circles. Hence, one has to resort to either an iterative method based on optimization with certain constraints for an approximate solution, or use image features first to initially place few cells suitably at locations with high feasibility and then fill residual gaps in an iterative manner. We chose the latter path for our method, as the FGR map (in figure 3(d)) provides rich clues to initially detect a few cells with high level of accuracy.

Our proposed method consists of two sub-processes (figure 4): *Phase-I* sub-process consists of steps, which enhance the foreground areas (FGR) followed by feature extraction, using the Distance Transform (DT) from edge map of FGR and ridge lines from the modulated DT (mDT) map. Peaks of mDT map and features on ridge lines correspond to candidate centers of detected objects of interest. *Phase-II* sub-process fills the cell areas (detected centers with edges) detected in *Phase-I* with the background color on FGR to obtain a residual map (RFM). Significant areas of the RFM are then iteratively filled using Hough Transforms on convex arc segments that were identified on the outer boundary edges of the objects in the RFM. The output of the cell center locations provided by the two phases are eventually merged using a union operation (spatial locations of very nearby cell centers are averaged) to create the final set of detected cell centers. Both these phases of the pipeline (unsupervised) process are described below.

## III. Phase-I: DT and Ridge based method for cell center detection

Salient regions of the green channel correspond to the nuclei of interneurons. Since these are our main objects of interest, we have used the green channel alone for further processing. Steps used in *Phase-I* of our algorithm (see figure 4) are:

- Normalization of the green channel
- Contrast enhancement to get FGR
- Edge detection on binary image of the FGR map
- Distance Transform (DT) on edge map
- Compute mDT map using DT and FGR
- Ridge line detection (computed only for large areas of FGR) on DT map
- Identify peaks on Ridge lines using mDT map
- Detect Bifurcation Points (BF) on the ridge lines
- Clustering of the peaks and BF to yield the list of putative centers of cell nuclei.

As a first step, the green channel has been enhanced using contrast enhancement (detailed in supplementary section S.1) to obtain the FGR. Figure 5(a) shows the FGR with the Ground Truth (manually annotated cell centers). The green channel has alone been thresholded to produce the binary image of FGR (also shown in figure S.1(e)). We then applied a Canny edge detection method [10] on this binary image of FGR to obtain an edge map. A distance transformation (DT) [36] on the edge map is then used as an intermediate feature which will eventually help to locate candidate positions of cells in FGR. The DT values corresponding to the background region of FGR are then suppressed to zero, leaving only the DT-map on pixels of FGR. Figure 5(b) shows binary (thresholded) map of FGR in figure 5(a), with the edge map overlayed and represented by the Magenta borders. Figure 5(c) shows the DT map extracted from the edge map of figure 5(b).

**Fig. 5:**
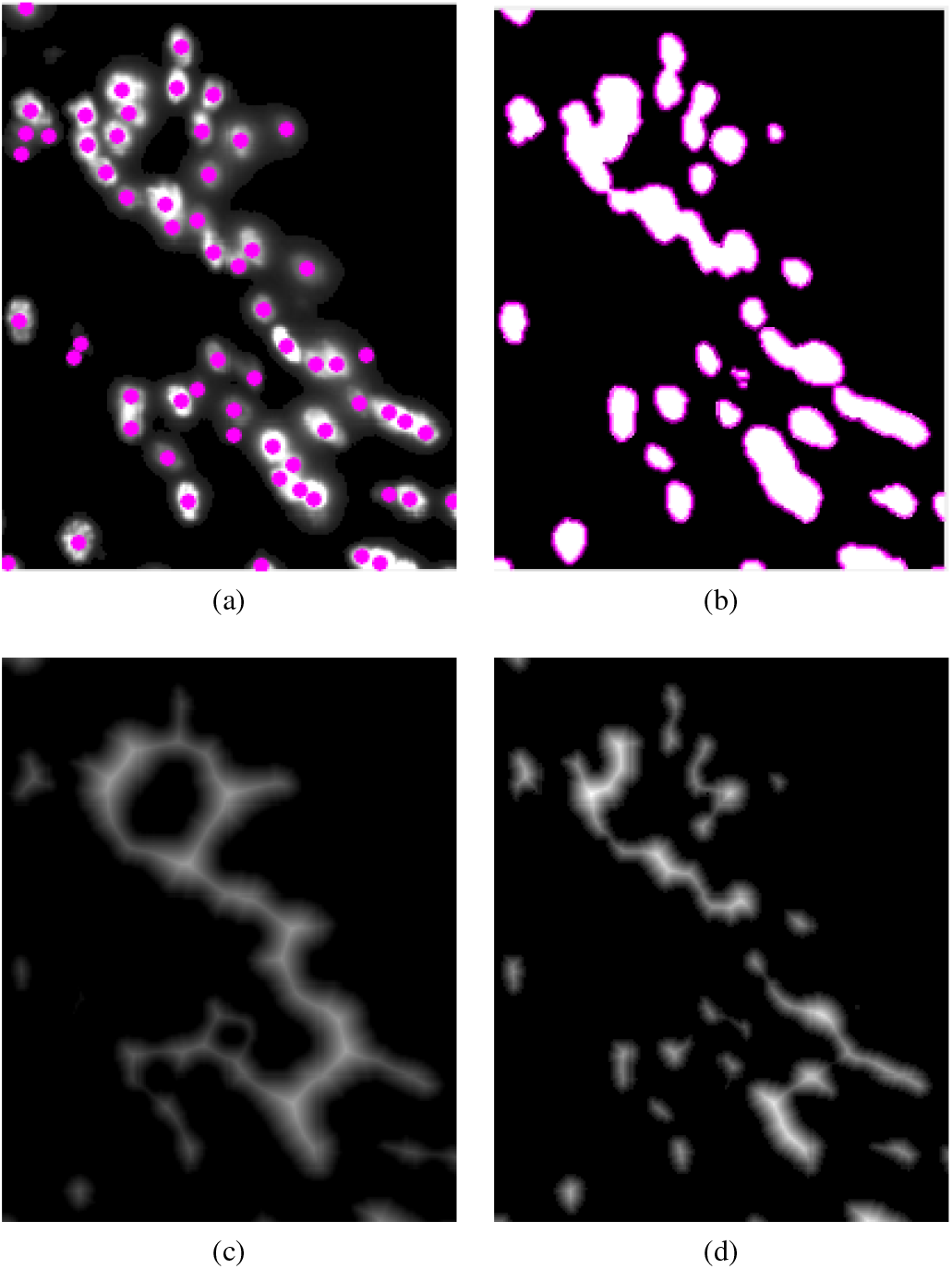
(a) Ground truth as manually annotated cell centers, on the green channel (only used in the figure) of the FGR shown in figure 3(d); (b) Edges marked in magenta over binary image (figure S.1(e)); (c) Distance Transform (DT) applied on image in figure S.1(e). DT is suppressed in the background region of FGR; (d) Modulated DT (mDT) map, obtained using equation (1) with FGR and DT map

The DT map [44] results in producing an intensity peak which corresponds to the centers of detected nuclei, and this intensity decreases the further we get away from the centers and thus the minimum intensity represent the edges. However, in cases of mild overlap, where several GFP nuclei overlap, DT map does not always exhibit sharp peaks, making it difficult to detect cells using only a local maxima. Hence we have used a modulated DT (mDT) map (figure 5(d)) obtained as:

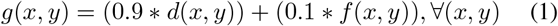

where, *g, d* and *f* represent mDT-map, DT value and FGR respectively, *x, y* represent integer pixel coordinates. The mDT map exhibits better localization of peaks as cell centers (as shown in figure 6(a)). Regional maxima in mDT map are thus detected and represent the putative centers of the GFP nuclei (figure 6(a)). The process described so far though works well in cases where the GFP nuclei are isolated and spatially distributed with mild overlap, however produces unsatisfactory results when few cells have strong (large) overlap in arbitrary spatial arrangements. As closely packed clusters of cells do not follow any well-defined structure or geometrical arrangement, expecting the center-line (as in MAT [13]) to be the loci of peaks would be an erroneous assumption. In such cases, the maxima in mDT map may be falsely detected to signify a single center when there are actually two or more cells present in the FGR (figure 5(a)) for that location. This process of detecting peaks in mDT, thus results in “misses” of cell centers for the complex cases of strong overlap of cells.

**Fig. 6:**
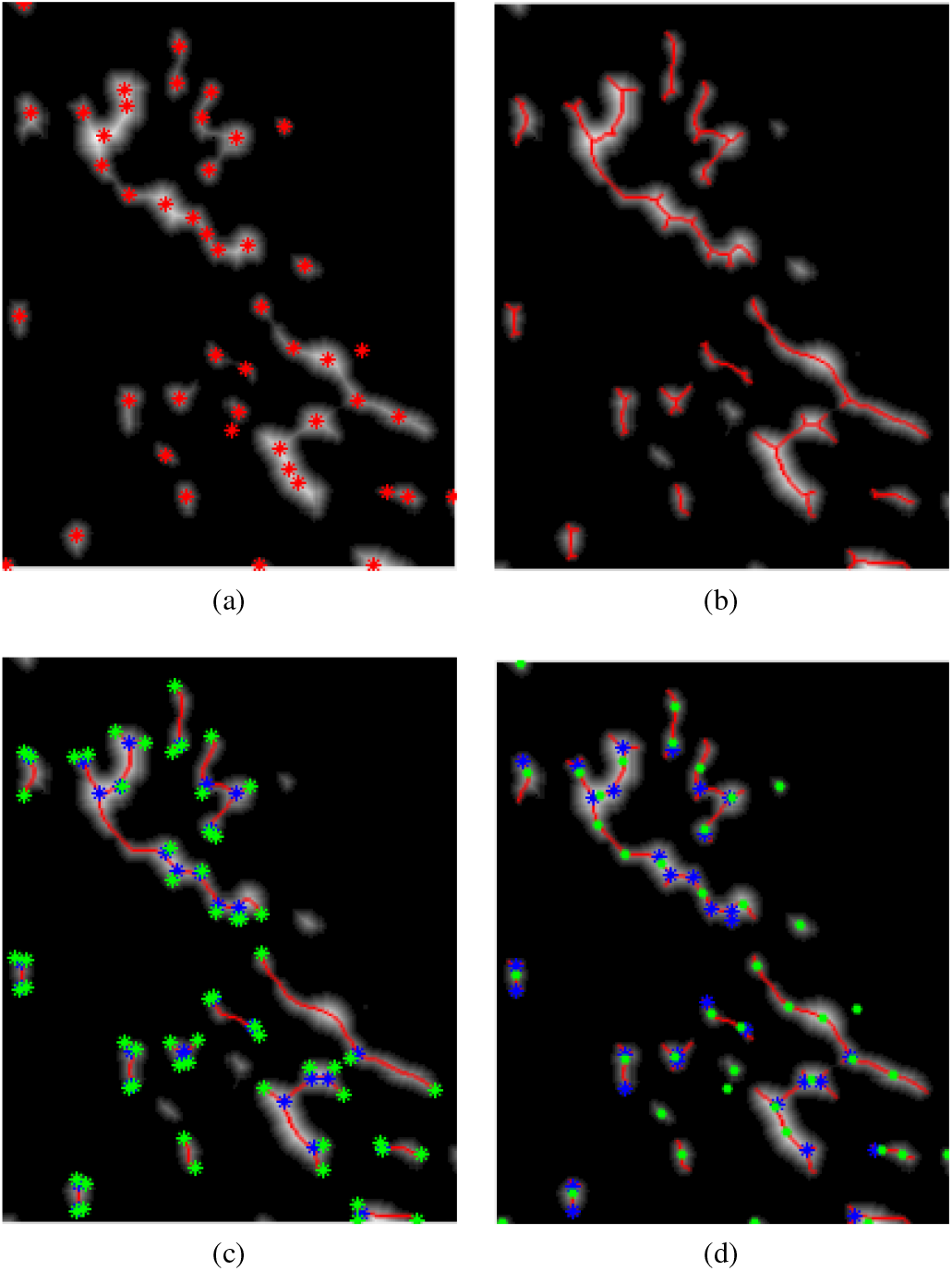
(a) Peaks detected on the mDT map as shown in figure 5(d); (b) the Ridge lines extracted from 5(d) using the “vessel” filter [23] algorithm; (c) Landmarks identified as Bifurcation points (BF) (in blue) and Ridge Endings (RE) (in green) detected on the Ridge lines as shown in (b); (d) Combination of peaks detected on mDT map (as in (a) but marked as green dots) and Bifurcation points (BF) (as in (c) marked as blue asterisk) marked on the Ridge lines in (b), detected as potential centers of the cell nuclei.

To overcome this problem, we have designed an algorithm that detects features within the ridge lines from mDT map (as shown in figure 6(b)) of only those putative cells whose area is greater than 1.5 times the average cell area (i.e., minimal and no overlap cells are not considered) and assigns the centers of the nuclei based on the location of these features on Ridge lines. This ridge detection process is similar to the “Vessels” Algorithm [23], that has been used earlier to detect blood vessels from magnetic resonance imaging (MRI) data. The features used in our analysis are the Bifurcation (BF) points and Ridge Endings (RE). RE appear near the border of the cells and only the Bifurcation Points (BF) correspond to potential cell centers. The BF were obtained as landmark features by traversing along the ridge lines (as in [49]). An example of those feature points detected on ridge lines is shown in figure 6(c). Traversing along the ridge lines, as shown in figure 6(c), produces an additional list of centers. These centers estimated on the ridge lines and those obtained from the peaks on mDT (as shown in figure 6(d)), are combined together (as a union of two sets, and merger of nearby peaks) to yield the final set of cell centers at end of *Phase-I* of processing. Figure 7(a) shows the results of intermediate level processing at the end of *Phase-I* (DT + Ridge-based method).

**Fig. 7:**
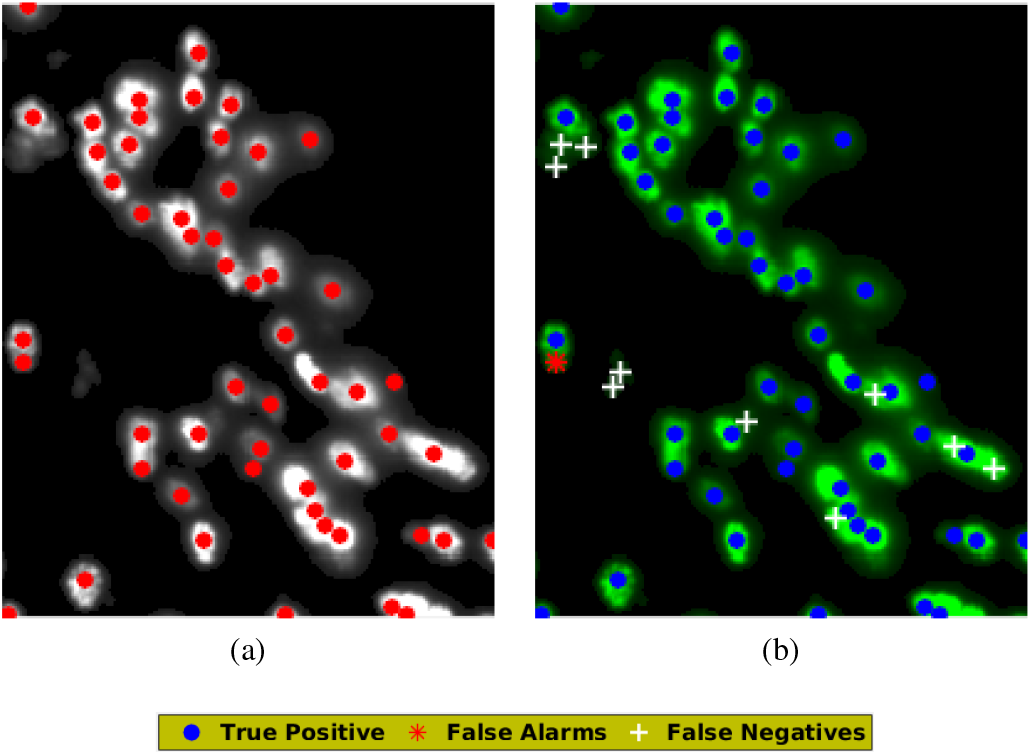
(a) Results as in figure 6(d) after merging and clustering the detected cell centers; (b) Evaluation of *Phase-I* results in (a), using true positives, false alarms and false negatives (best viewed in color).

## IV. Phase-II: Convex-Arc Based Iterative Method for missing cell detection

Results of *Phase-I*, though fairly accurate, did not yield satisfactory results in cases of strong (large) overlap of cells. An example of this failure is highlighted by figure 7(b) where true detections, misses and false alarms of *Phase-I* are shown alongside true centers as annotated by a master annotator.

To further enhance the efficiency of the algorithm, we have introduced an iterative process (algorithm 1) of cell filling using boundary arcs on a residual foreground layer (FGR in figure 5(a)). This process is based on the hypothesis of topological/geometric constraint: in case of strong overlap, cells at the periphery of the region of overlap produce the boundary (edge) layer of FGR. Those in the inner cordon fill up the gap to produce prominent green fluorescence within the FGR. To efficiently detect the overlapping cells, an iterative process is designed to detect overlapping cells in FGR, starting with boundary layer, which is described below as algorithm 1.

**Algorithm 1:**
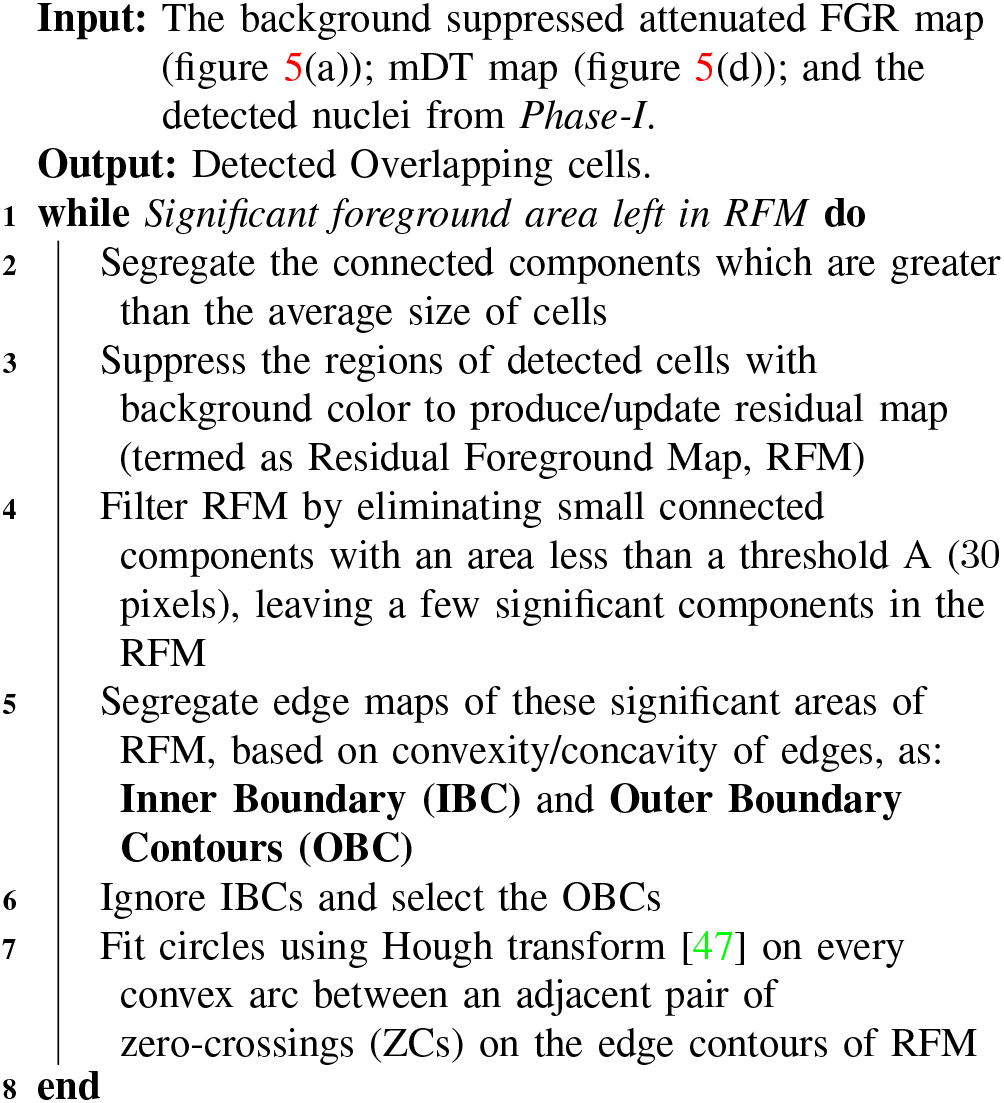
Detecting Overlapping cells at *Phase-II*.

This unsupervised process iteratively identifies undetected cells over strong overlapped regions; specifically over significant RFM areas, which are missed at the **Phase-I** of processing. A flowchart describing algorithm 1 along with a synthetic example is discussed in detail in the Supplementary section S.2.

Figure 8 demonstrates the application of the iterative arc based method for missing cell detection in the given example from figure 5(a). Figure 8(a) shows the peaks detected at *Phase-I*. Figure 8(b) gives residual map (RFM) obtained after suppressing the detected cell regions in Figure 8(a) with the background color. Figure 8(c) shows Inner boundary contours (IBCs) in green and Outer boundary contours (OBCs) in blue on the significant RFM after filtering out the small connected components from that in Figure 8(b). IBCs are eliminated and convex arcs are fitted (using Hough Transform) with circles on convex OBCs. Figure 8(d) shows the centers of the new cells detected at *Phase-II* of processing. The final list of cell centers is obtained by a merging process of the two lists of cell centers detected from Phase I: DT + Ridge-based peak feature detection (e.g. Figure 8(a)) and Phase II: Arc based iterative filling (e.g. Figure 8(d)) methods. If two or more distinct cell centers detected at *Phase-I* and *Phase-II* stages fall within a small neighborhood (*dc* < 2 ∗ *RC, RC* = average radius of a cell), they are merged and the average of the locations of the cell centers replaces the set of nearby cells. Figure 9 gives an illustration of the results of this process of merging and detecting new cells in residual foreground layer. Supplementary section S.3 contains the sequence of intermediate results of different steps of processing in sub-processes *Phase-I* and *Phase-II*. This section also contains the table of all heuristic parameters used in our model for cell detection.

**Fig. 8:**
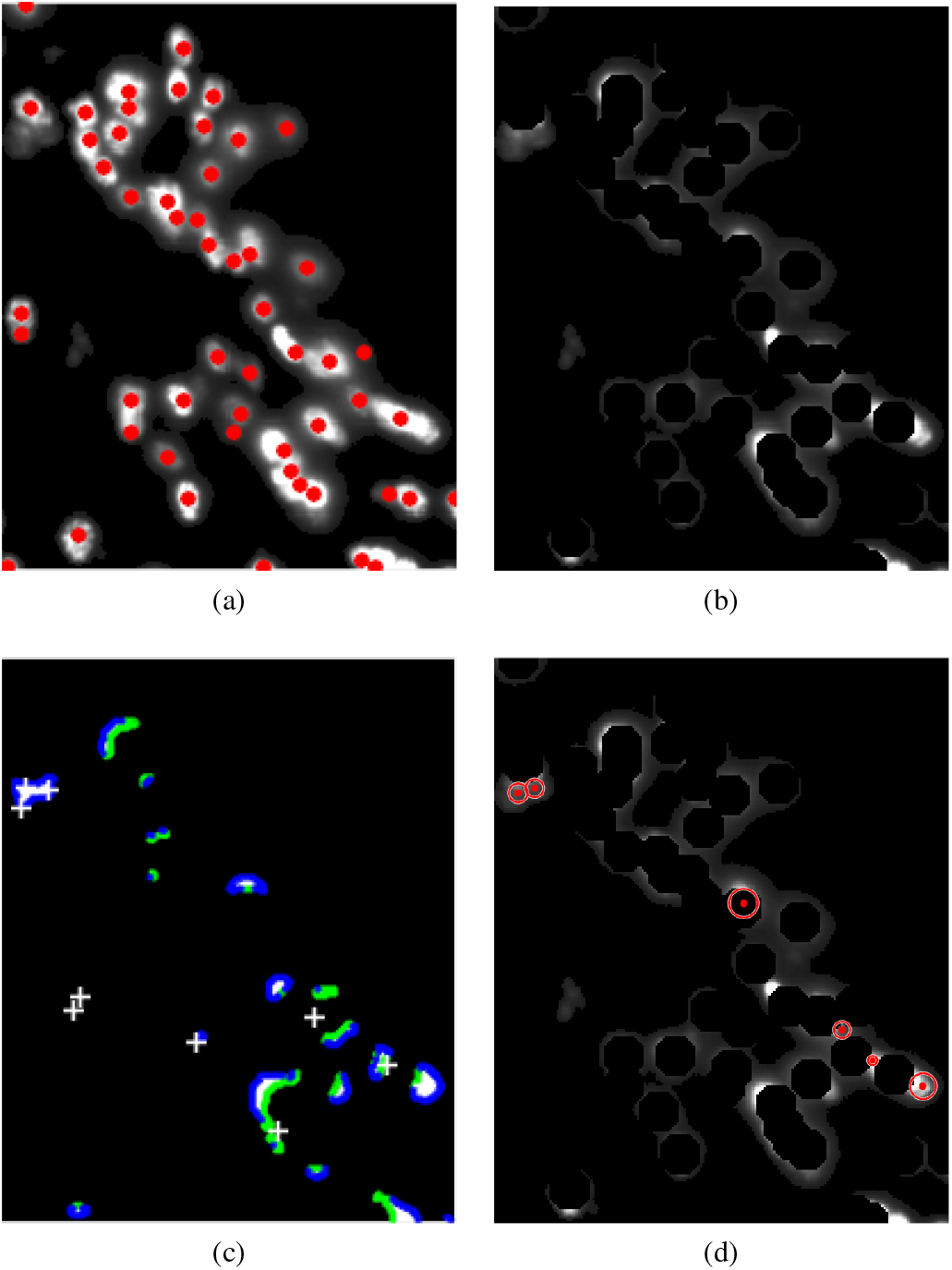
Results of application of *Phase II* process on the example section used in figures 3(d), 7 containing strong overlap of GFP nuclei; (a) Centers detected by *Phase-I* of processing on FGR, as given in Figure 7(a); (b) RFM (Residual foreground map) obtained after first iteration, from (a); (c) IBC in green and OBC in blue with the missed cells marked on the binarized RFM in (b). Contours are obtained from (b) using Canny’s [10] edge detection algorithm; and (d) cell centers detected by fitting circles using Hough Transform only on convex OBCs (figure best viewed in color).

**Fig. 9:**
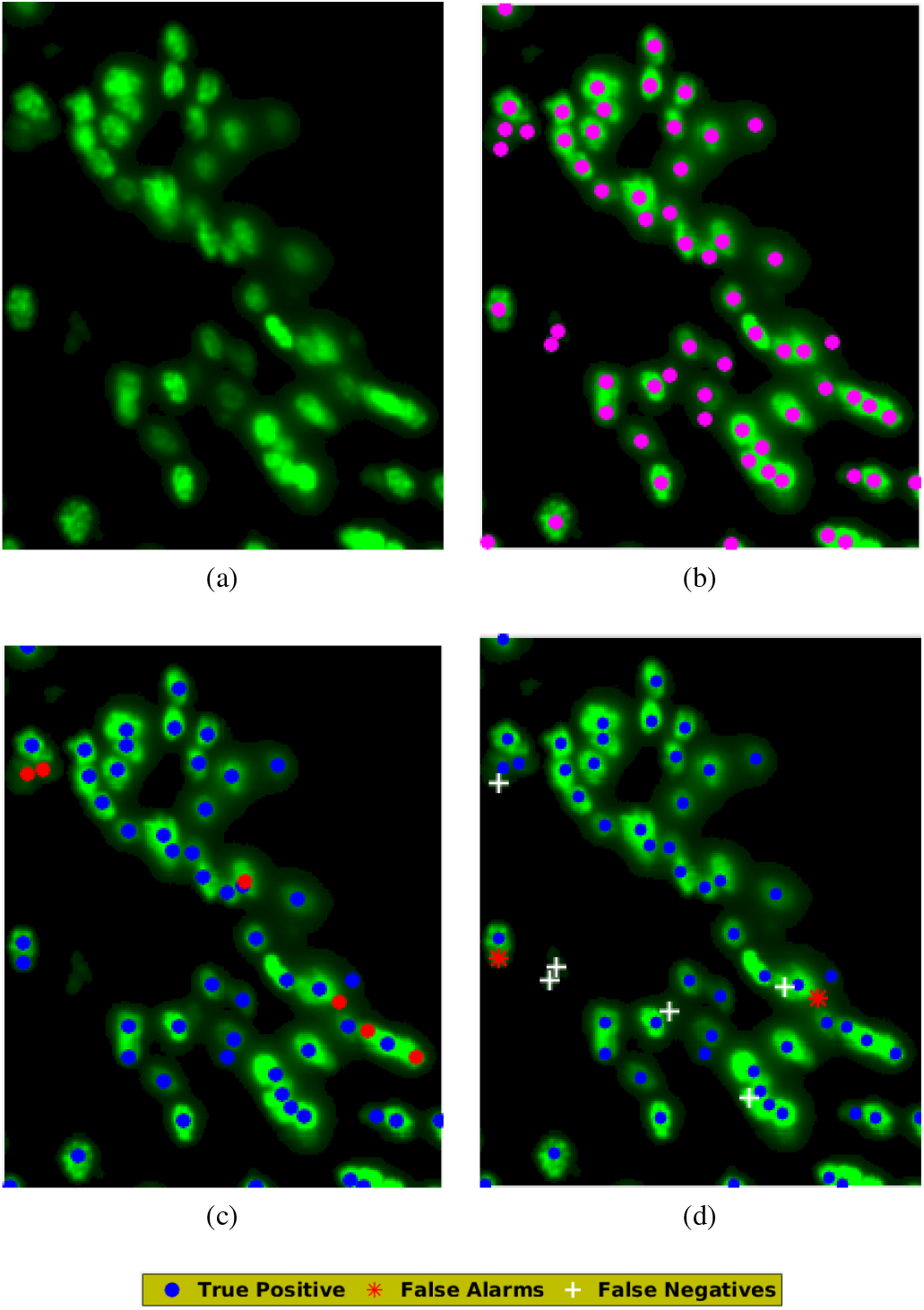
(a) RGB image of FGR (with non-green channels suppressed) as shown in Figure 3(d); (b) The ground truth obtained by manual annotation on (a); (c) Cell centers (GFP nuclei) obtained after merging centers obtained from *Phases I* and *II* of processing on (a), shown in blue and red dots respectively; and (d) gives true positives, false alarms and false negatives (best viewed in color).

## V. Results and Performance Analysis

We have evaluated our algorithm on gigapixel resolution brain section images of two mouse brains. The data [1] was manually annotated by a set of annotators (*n* = 12; 10 naive and 2 master annotators) to obtain the groundtruth. Each gigapixel resolution image was split into tiles of 500 × 500 pixels for easier quantification by the human annotators. Each annotator was also given the classification results of the DT and ridge based method (*Phase-I*), and then allowed to modify, insert or delete the existing cell centers. The output of the *Phase-I* was provided to reduce the burden of identifying all isolated cell locations by visual search. This did not bias their judgment in the complex regions of overlapped cells. Annotators easily modified (delete, insert, change location) the detected cells using a GUI (see snapshots in section S.7 of the Supplementary Document). Providing manual annotators the output of a first-pass algorithm is “industry standard” in related problems (*e.g*. in EM for image segmentation). The annotation labels were cross-verified by a master annotator, which gave us confidence of its accuracy levels.

Figure 10 shows typical examples of cell detection for selected parts of mouse brain scans obtained using our proposed method. The proposed algorithm takes the cell centers detected from two phases, *Phase-I*: DT + Ridge-based; and *Phase-II*: Arc-based methods, and combines them using an iterative combination strategy. True positives, false alarms and false negatives are marked with blue, red and white color markers respectively in the outputs. *Phase-I* sub-process is sufficient to detect all cells accurately in sparse and minimal overlap regions. The Phase II sub-process requires 2 – 3 iterations to converge, typically in cases of strong overlapped cells. The details of training the supervised architectures are given in the next subsection, and are elaborately discussed in the Supplementary section S.4.

**Fig. 10:**
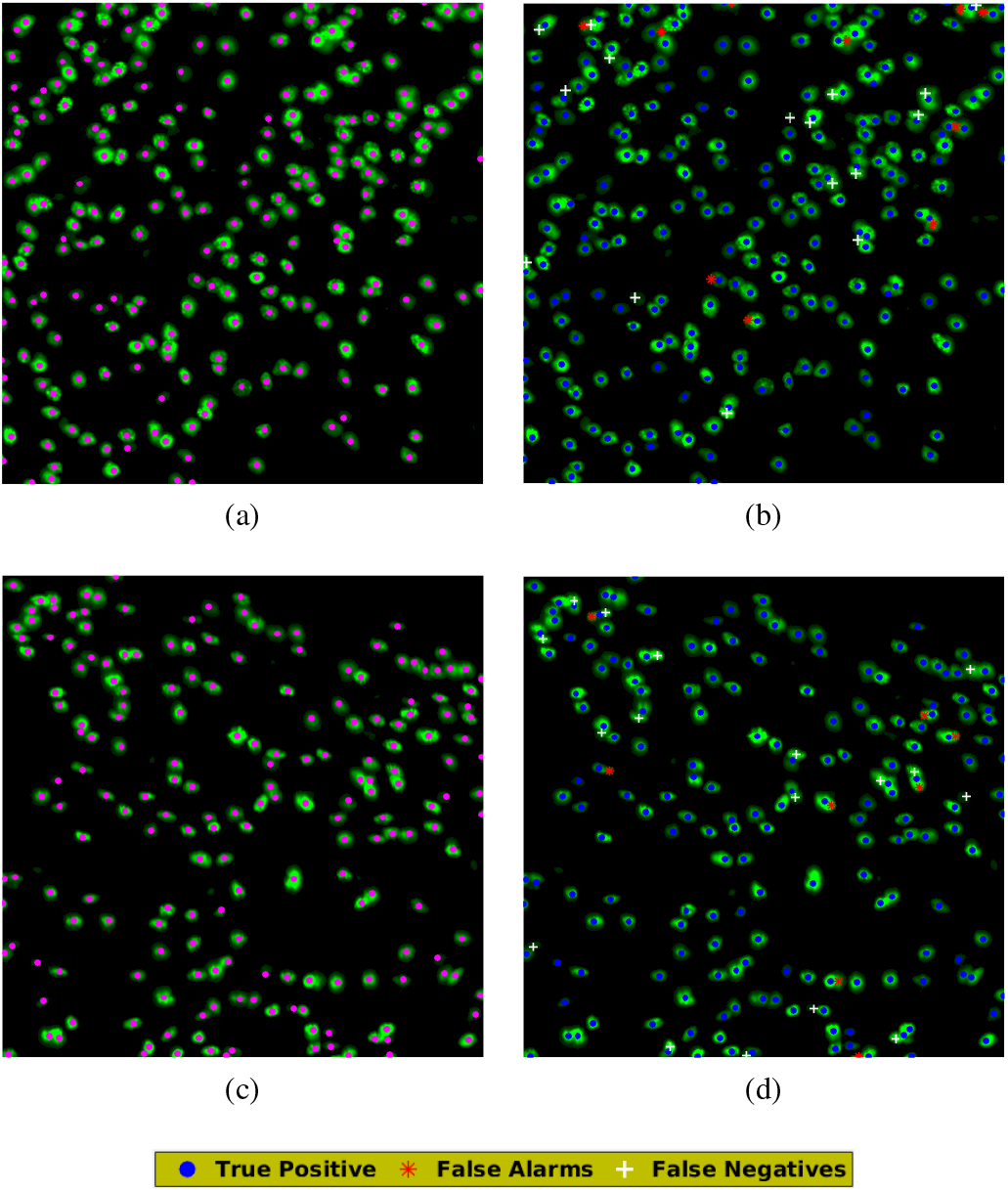
Result of cell detection process using proposed method (figure 4); (a) and (c) show two different regions with strong and minimal overlap from Hua-166 and Hua-167 scans respectively with ground truth; (b) and (d) are results evaluated from our proposed method corresponding to scans in (a) and (c) respectively (best viewed in color).

### A. Evaluation of the proposed algorithm

The manually annotated samples were used as groundtruth to evaluate the performance of the algorithm. In addition, the performance of our proposed method was compared with other methods that can potentially be applied to solve the same problem. These are two supervised shallow learning methods: (a) SVM [16], [41] (using Gaussian Kernel); and (b) HAAR + Adaboost [57]; (c) cell detection based on the method used by Al-kofahi *et al*. [3]; two DL techniques: (d) Faster R-CNN [50] and (e) SegNet [6] and (f) *Phase-I* of our proposed approach (for comparing with *Phase-II*). For the supervised approaches, we have used half of the annotated data for training and the remainder of the data for testing. A Precision-Recall metric was used as the performance criteria [42]. The cell-centers that were detected within a certain distance (DR < 5 pixels) of true location in groundtruth were considered as to belonging to the same cell. Cell-centers of positively classified windows are considered as detected cell centers. The details of training the supervised architectures are elaborately discussed in the Supplementary document (section S.4 for Faster R-CNN and SegNet).

**TABLE I:**
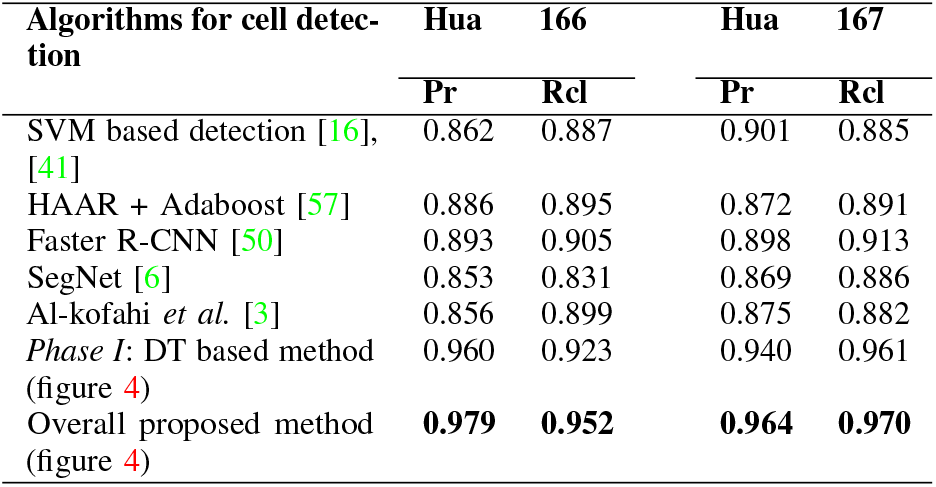
Performance of our algorithm averaged over two sets of mouse brains (selected 158 scans of *Hua-166* and all 244 scans of *Hua-167*, with 18*K* × 24*K* pixels each). Our approach provides the best precision and recall over supervised machine learning (shallow learning with 5-fold validation and deep learning methods) approaches. ***Pr*** - Precision; ***Rcl*** - Recall.

We have used a tensor flow based implementation of Faster R-CNN [50] for our experiments. Faster R-CNN [50] performs a bounding box regression on the objects in an image for detection. We have used equal number of samples for training and testing. Since we have only one object of interest i.e. GFP labeled neuron in our data, we provide a bounding box of size 25 × 25 pixels (derived empirically) around each cell. Faster R-CNN [50] uses the co-ordinates of the positive samples as the anchor boxes. Centroids of the bounding boxes obtained as output are considered as detected cell centers. We have utilized a SegNet [6] model that has been trained as per the code provided by the authors. Training samples (only positive ones) in SegNet were generated by delineating circular patches around manually annotated cell centers. The inputs for training were circular patches of 4 pixels radii around the manually annotated cell-centers. The trained SegNet map produces outputs as circular patches (sometimes overlapping with each other) on the test image, whose centers can be considered as putative cell-centers. Details of training are given in Supplementary section S.4. Our method has also been compared with that of Al-kofahi *et al*. [3], which, like our proposed model, is also an unsupervised method used for cell nuclei detection and segmentation. It initially extracts the foreground region using graph-cuts-based binarization, following which a multiscale Laplacian-of-Gaussian filtering combined with distance-map-based adaptive scale selection is used to detect the seed points for segmentation. It then detects nuclei through graph-cuts-based segmentation along with alpha expansions and graph coloring.

**TABLE II:** Performance of proposed unsupervised method evaluated separately on the different sections of the brain, with strong overlap and minimal overlap of cells, for the proposed method; **Pr** - Precision; **Rcl** - Recall.

| Performance our proposed method on Cell distributions in Mouse brain scans | Hua-166 |  | Hua-167 |  |
| --- | --- | --- | --- | --- |
|  | Pr | Rcl | Pr | Rcl |
| Strong Overlap of multiple cells, in parts of $\sim 5$ scans/brain (Overall) | 0.95 | 0.94 | 0.96 | 0.968 |
| Strong Overlap of multiple cells, in parts of $\sim 5$ scans/brain (Phase-I) | 0.92 | 0.89 | 0.90 | 0.88 |
| Minimal overlapped Cells, in parts of $\sim 20$ scans/brain (Overall) | 0.96 | 0.95 | 0.97 | 0.97 |
| Minimal overlapped Cells, in parts of $\sim 20$ scans/brain (Phase-I) | 0.93 | 0.87 | 0.90 | 0.88 |
| Sparse Cells, in abundance $\sim 100$ scans/brain (overall/Phase-II) | 0.978 | 0.96 | 0.98 | 0.98 |

Comparative analysis of our algorithm with the other models are tabulated in table I. Our iterative approach exhibits high precision and recall values of 0.972 and 0.961 (average over complete dataset of one mouse brain and a selected portion of another) when evaluated using the manually annotated ground truth, indicating the close to human efficiency of the algorithm. The supervised methods of cell detection based on SVM and Faster R-CNN based deep learning did not perform better than our proposed unsupervised approach. Details and justification of using a Non-Maximum Suppression (NMS) to obtain the results of all supervised methods are given in Supplementary section S.5.

**TABLE III:**
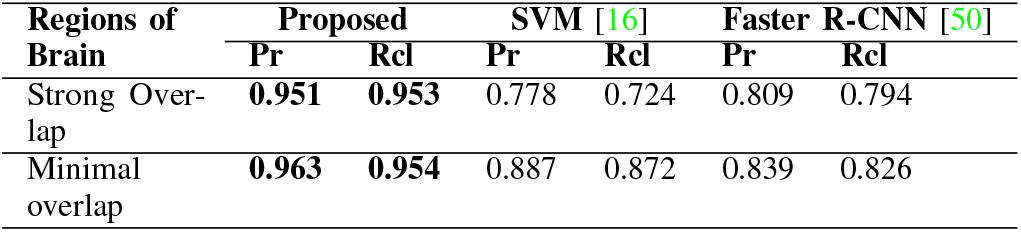
Comparative Performance over the strong and minimal overlap regions of cells (from ≃ 400 scans of two mouse brains), for the proposed, SVM and Faster R-CNN methods.

| Regions of Brain | Proposed |  | SVM [16] |  | Faster R-CNN [50] |  |
| --- | --- | --- | --- | --- | --- | --- |
|  | Pr | Rcl | Pr | Rcl | Pr | Rcl |
| Strong Overlap | **0.951** | **0.953** | 0.778 | 0.724 | 0.809 | 0.794 |
| Minimal overlap | **0.963** | **0.954** | 0.887 | 0.872 | 0.839 | 0.826 |

Our proposed unsupervised iterative method of cell detection, provide near-ideal performance in most of the cases, with exceptions that can be attributed to the presence of noise in the dataset. In order to demonstrate the effectiveness of the algorithm over wide distribution of cell nuclei throughout the brain, we have separated the results over strong overlap and minimal overlap and sparse regions within cells of the brain, and results for both sub-processes are given in table II. As expected, the algorithm performs the best in the sparse regions with marginal degradation in performance in the minimal overlap and strong overlapping areas. Table III shows the comparative performance, on the separated data sets of overlapped, sparse and minimal overlap regions of the brain, of our proposed method compared to that of SVM [16] and Faster R-CNN [50]. Additional results (with examples) of cell detection of the competing supervised methods used for performance analysis are given in Supplementary section S.6.

The proposed iterative approach (*Phase II*) converges in typically 2 – 3 iterations for the complex overlap cases. For the example shown in figure 9, the algorithm took *1327 ms* for complete evaluation compared with *795 ms* for the *Phase-I* stage on an Intel core-i7 4820K processor, with 64 GB RAM. The average time taken by the proposed method for a single brain section of 18*K* × 24*K* pixels on an Intel core-i7 4820K processor based CPU with 64 GB RAM is 4 – 5 *mins*. Despite having doubled the number of training samples, the competing supervised methods did not exhibit better performance, indicating that evaluation of the model is sound. A 5-fold study has been performed for shallow ML models (5 different combinations of randomly selected sample sets to form the training and test partitions). For deep-CNN we have done a 2-fold study; and our unsupervised method does not require training samples; all samples are used for testing, a few are used to empirically obtain the optimal parameter values.

## VI. Discussion

Our approach allows for robust and efficient estimation of the centers of GFP tagged cell nuclei with relatively high precision and recall. Our unsupervised and iterative algorithm based on Ridges and distance transform [44] combined with arc-based region filling provides the best precision value amongst the different approaches used to solve the detection problem. It is important to note that all the methods (including shallow and deep learning methods) also perform well in cases where there is marginal or no overlap (sparse distribution) between the GFP tagged interneurons. Our algorithm approaches near ideal performances in these cases, with the precision-recall values ≃ 0.97 and ≃ 0.96 respectively. The cases where the other models fail are the complex case of strong overlap of cells, where our method gave the best result with a overall precision of 0.972 and recall of 0.961. The drawback with our algorithm is the relatively longer evaluation time frame which is directly proportional to the complexity of the image to be evaluated. Our proposed algorithm is completely automated and unsupervised in its application. We have applied this algorithm to analyze brains and a web implementation of the iterative algorithm has been implemented at the mouse brain architecture portal at the Cold Spring Harbor Lab (CSHL) [1] for analyzing cell type data.

It was surprising to note that standard supervised classifiers, e.g. SVM, ADABOOST, etc. and state-of-the-art DL based detection techniques (Faster R-CNN and SegNET) did not perform satisfactorily when compared to our method. We believe that this may be because of our method of treating strong overlap decomposition explicitly which improves our performance to ~ 97%, the precision levels needed for real world application. The performance of the ML models using our method is comparable with a previous study on brains [28] where similar methodology of GFP tagging in mice were used and the detection was performed with CNN in order to achieve 90% performance. The difficulties for the network may have been due to the degree of overlap of objects in the image, which, in turn, depends on cell densities as well as microscopic methodology.

## VII. Conclusion

We have developed an iterative approach (unsupervised) for efficiently detecting GFP tagged cell centers of GABAergic neurons in the mouse brain. Our results demonstrate that our algorithm, which is based on classical machine vision methods substantially outperform published CNN-based methods. The detection problem that we have solved is of practical interest to neuroscientists. The classical Machine Vision based method we have designed, directly solves the overlap decomposition problem with high precision and recall values. Recent methods of supervised object detection do not provide the level of performance required particularly in cases of strong overlap of cells, as the foreground blobs appear with a wide variety of non-unique two-dimensional patterns that are of different shapes, sizes and structures (due to varied spatial arrangements of cells in strong overlap), making the training data unreliable for efficient learning. Our unsupervised approach provides a much superior performance than recent supervised object detection algorithms such as ADABOOST [57], SVM [16], [41], SegNet [6] and Faster R-CNN [50]. Previously published results [28] on similar problems only reach 90% performance level (in contrast with > 97% of ours). The failure of both the deep and shallow nets to reach > 90% accuracy levels, is mainly attributable to the difficulties of accurately detecting overlapped nuclei. While our method may not generalize to other unrelated problems, it addresses an important problem in the growing field of computational neuroanatomy of brains. CNNs and SVMS are very powerful ML architectures but need further refinements to solve every real-world problem. In our use case the structure of the problem demands a different architecture.

With an ever increasing dataset and information about the neurons and their connectivity, it becomes important to develop automated unsupervised algorithms for efficient data analysis of relevant information [5]. We have provided one approach towards this goal which helps neuroscientists to achieve few targets with greater confidence.

## VIII. Acknowledgments

The work was supported by the HN Mahabala chair professorship at Indian Institute of Technology, Madras, The Crick-Clay Professorship at the Cold Spring Harbor Laboratory to Dr. P. Mitra. The authors also acknowledge the support from the Mathers foundation, NIDA grant (5R01DA036400-03) and NSF EAGER grant (1450957).

## S.1. Description of Contrast Enhancement sub-process in *Phase-I*

This section explains the process of obtaining foreground region (FGR) from a brain section. Let *S_G_* denote the green channel of a brain section *S*. As a first step, the green channel *S_G_* has been normalized to get the normalized green channel *I_G_* (as shown in figure S.1(b)). If *L* is the number of gray levels in *S_G_*, then the normalization operation can be expressed as:

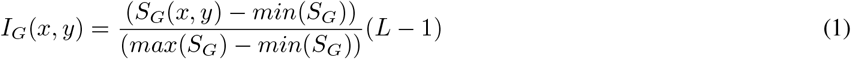

where *min*(*S_G_*) and *max*(*S_G_*) denote the smallest and the largest intensity values in *S_G_* respectively and *x* and *y* denote the pixel coordinate values. To enhance the perceptibility and improve the contrast of *I_G_*, a simple contrast stretching is performed resulting in a visually enhanced and a better contrast image *I_CE_* (as shown in figure S.1(c)):

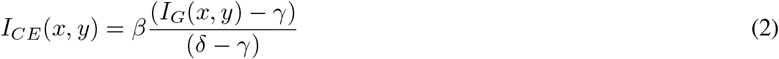

where *β = L* – 1. *γ* and *δ* represent the lowest and highest pixel intensity parameters of *I_G_* that need to be mapped to the new range in *I_CE_*. Since any real world image is susceptible to noise, choosing *γ* and *δ* as the lowest and highest pixel intensities of *I_G_* might have a horrendous impact on the stretching. So the bottom and top 1% (denoted as *th_l_* = 0.01 and *th_h_* = 0.99) of the brightest and darkest pixels have not been taken into consideration. Also, the intensity values in the image *I_G_* are clamped to get a new image *J* which is used for contrast enhancement to obtain *I_CE_* as follows:

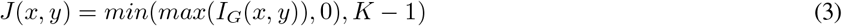

where *K* = 256. This can also be expressed as:

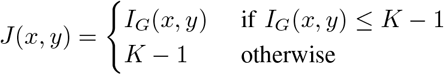

**Fig. S.1:**
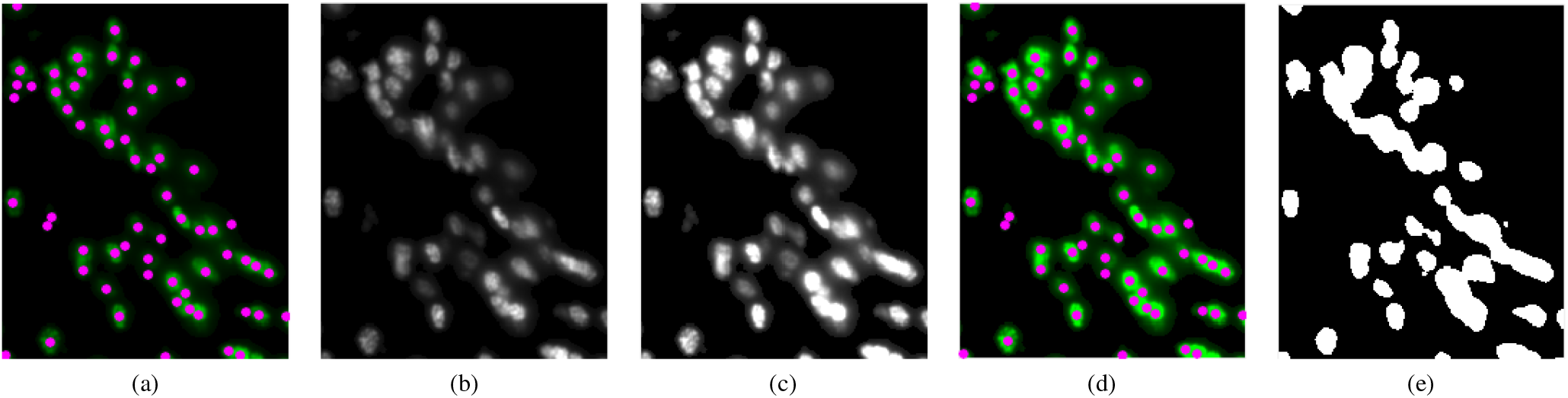
(a) Ground truth (marked as magenta dots) shown on the sample in Figure 3(b) (in the main manuscript); (b) the green channel of Figure 3(b); (c) Contrast enhanced image (FGR) of (b) obtained using equations 1-5; (d) Ground Truth on RGB image of (c) (same as Figure 3(d)) with non-green channels suppressed; (e) Binary image of (c), also used in Figure 5(b). (Figure best viewed in color)

Let *P_c_* denote the cumulative distribution function (CDF) obtained from the histogram of *J. γ* and *δ* are now obtained as:

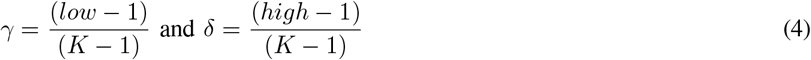

where

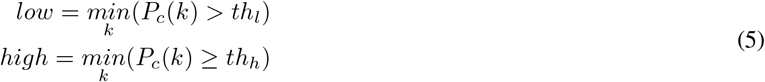

where *k* = 0, 1,…, *K* − 1.

## S.2. Description of *Phase-II* sub-process using a synthetic example

Various stages of the iterative process within the *Phase-II* sub-process (as described in algorithm 1 in the main manuscript) are elaborated further in this section. Figure S.2 is a flowchart describing the same. figure S.3(a) shows a synthetic example of 4 overlapping cells, given as input to *Phase-I* of processing. Results of various intermediate stages of the entire process of cell detection are shown in the rest of the figure S.3. The detail steps of the iterative process (*Phase-II*) to detect strong overlap of cells, using the synthetic example (as shown in figure S.3) are detailed below figure S.3.

**Fig. S.2:**
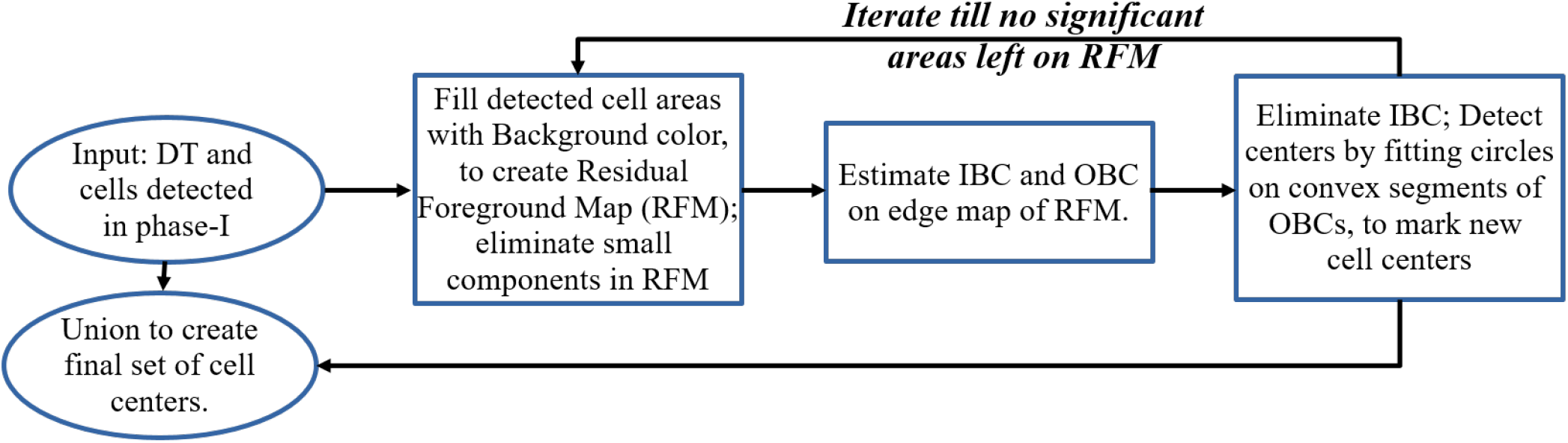
Iterative arc based method for cell detection. This phase uses results from *Phase-I* of the algorithm as inputs and the final result of nuclei centers is the combination of the output from this phase and that from *Phase-I*.

**Fig. S.3:**
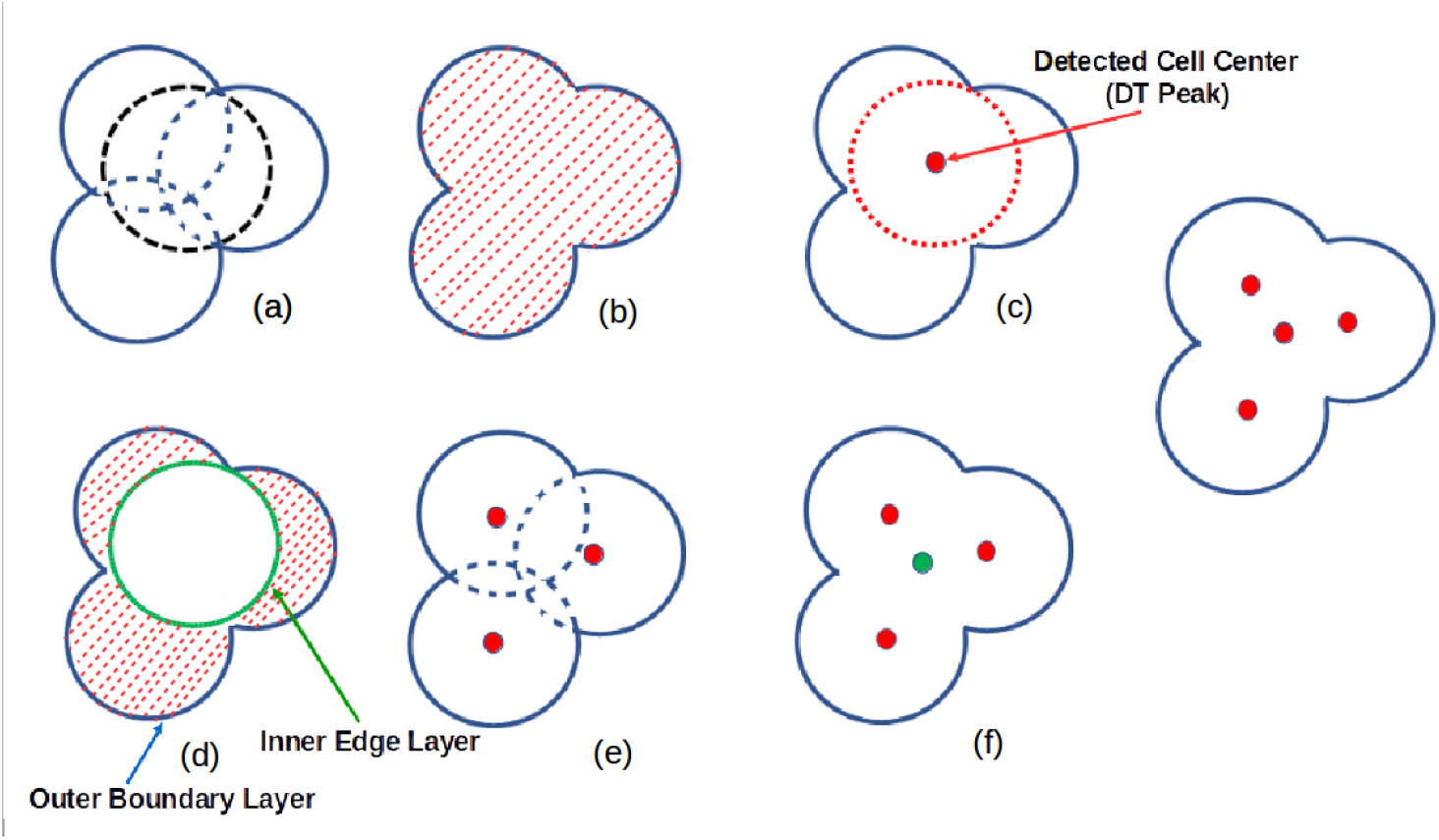
A synthetic example to illustrate the processing steps of *Phase II* of the algorithm: (a) Simulated overlapping cell regions, (b) Edge map of the overlapping cell regions with FGR shaded; (c) detected cell center (using mDT Peak) and cell region, (d) Residual area, RFM, after suppressing the cell regions (detected in (c)); also, Inner Boundary Contour (IBC) marked in green and Outer Boundary Contour (OBC) marked in blue, (e) Circle fitted on the convex arcs or OBC using Hough Transform (HT), followed by extension to the rest of the circles shown in dashed curves and centers; (f) new cells detected in red along with center detected in green from (c); and (g) final set of detected centers on the synthetic example of strong overlap of cells in (a).

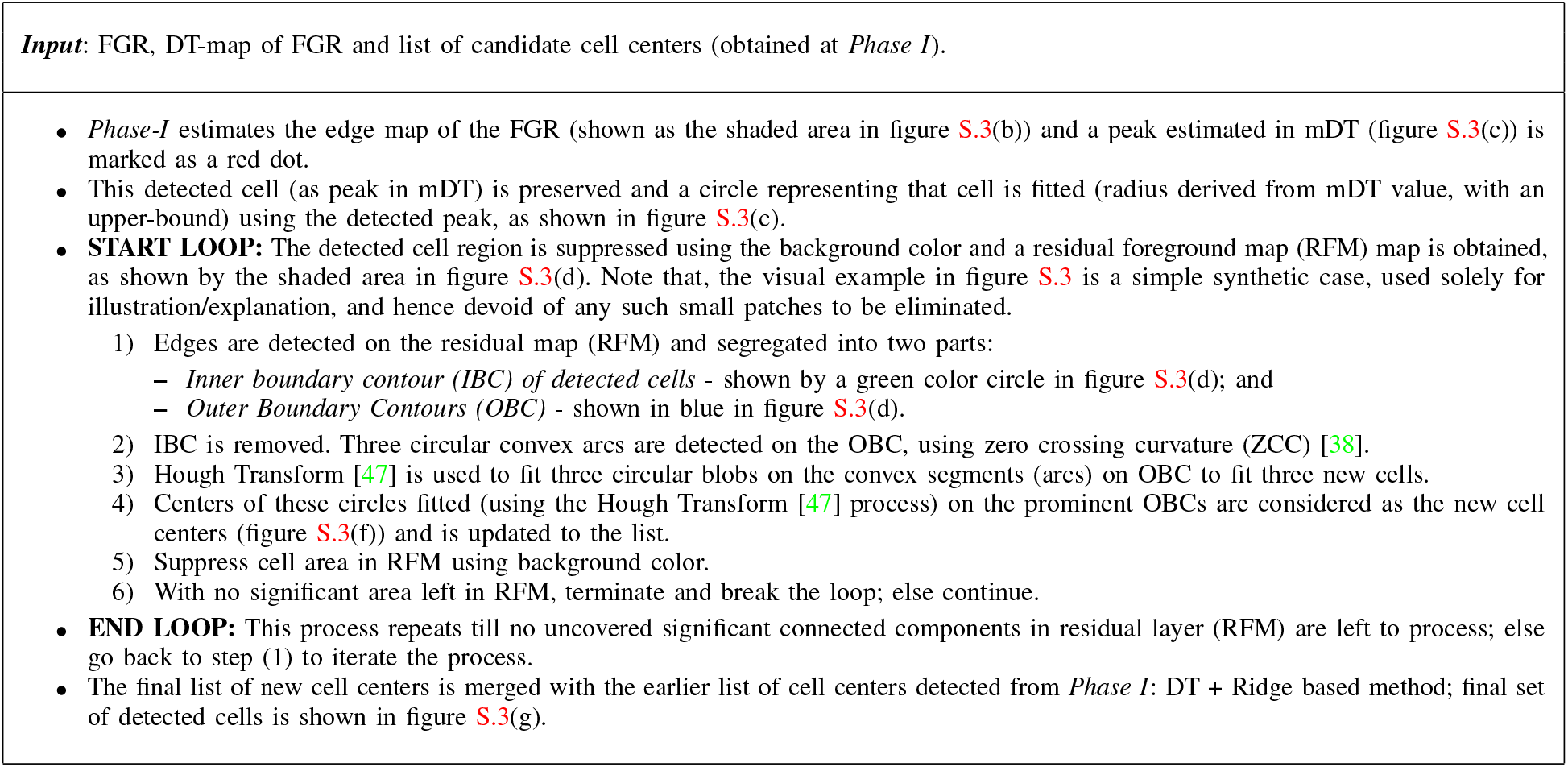
Detail of iterative steps in sub-process *Phase-II*

## S.3. Intermediate results of the diffe rent steps in the sub-processes: *Phase-I* and *Phase-II*

This supplementary section shows the intermediate results of both the sub-processes in the proposed pipeline (figure 4) for cell detection. It also illustrates the optimal parameter values used in the algorithm as a table. Figure S.4 illustrates the results of the intermediate steps in *Phase-I* sub-process, for the example given in Figure 3(d) in main manuscript. Following the steps given in section III of manuscript, results of eight intermediate steps are shown.

**Fig. S.4:**
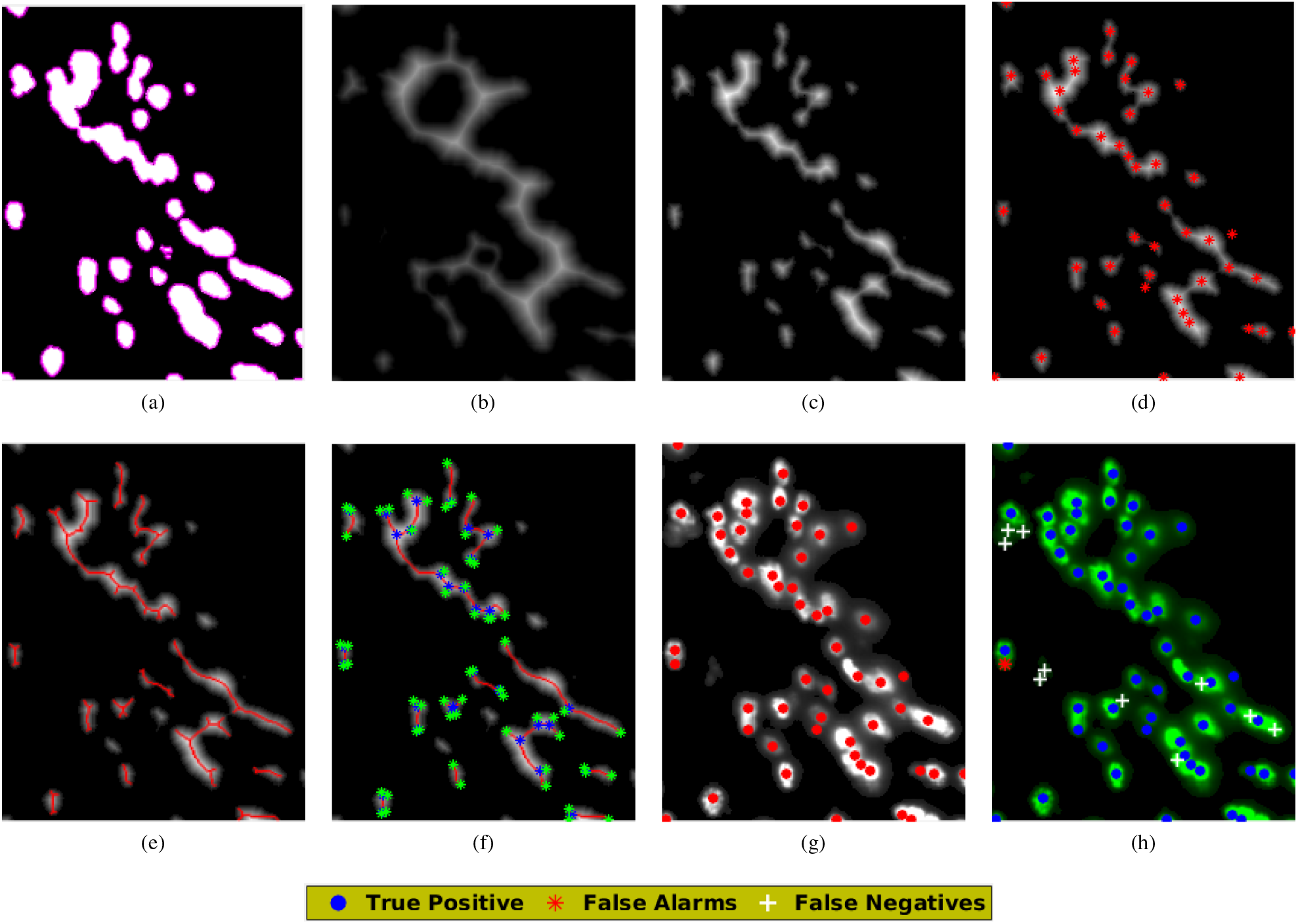
(a) binary image (of the example given in S.1(a)) overlayed with the edge map (in magenta) obtained using Canny edge detection algorithm (same as in Figure 5(b)); (b) Distance Transform (DT) on edge obtained from S.1(e); (c) Modulated DT (mDT) map, obtained using equation (1) (in the main manuscript) with FGR and DT map; (d) Intensity peaks (red dots) marked on mDT map in (c); (e) the Ridge lines extracted from (c) using the “vessel” filter [23] algorithm; (f) Landmarks identified as Bifurcation points (BF) (in blue) and Ridge Endings (RE) (in green) marked on the Ridge lines as shown in (e); (g) Final set of Cell Centers obtained by merging (union and clustering) nearby cell center positions obtained in (d) and Bifurcation points (BF) in (f); (h) Evaluation of the results of *Phase-I* sub-process compared with the ground truth, showing the true positives, false alarms and false negatives marked on enhanced image in Figure 3(d) (of main manuscript) (figure best viewed in color).

### A. Heuristic parameters used in the model for cell detection

**TABLE S.1:** Values of heuristic parameters used in the proposed algorithm (refer sections III & IV of the main manuscript). All values are obtained empirically to be optimal.

| Params | Phase | Details & Remarks | Value |
| --- | --- | --- | --- |
| *th* | I | Threshold used for binarizing FGR | 0.25 |
| *ar* | I | Area threshold for ignoring smaller connected components of FGR for Ridge estimation | 54 pixels |
| *dc* | I | Minimal distance for Merging Centers of cells | 12 pixels |
| *CD* | II | Average Diameter of cells (exact value obtained using DT) | 4–7 pixels |
| *A* | II | Area threshold for detecting significant connected components in RFM; Phase II - Arc Based Iteration stage | 30 pixels |
| *DR* | I & II | Evaluation Process | 5 pixels |
| *W* | - | Window size for NMS (suppression of non-maximas based on confidence values) of Faster R-CNN, SVM & ADABOOST methods | 11 pixels |

Figure S.5 shows the results of the intermediate steps in *Phase-II* sub-process, for the example given in Figure 3(d) of the main manuscript. Following the steps given in section IV of manuscript, results of eight intermediate steps are shown.

**Fig. S.5:**
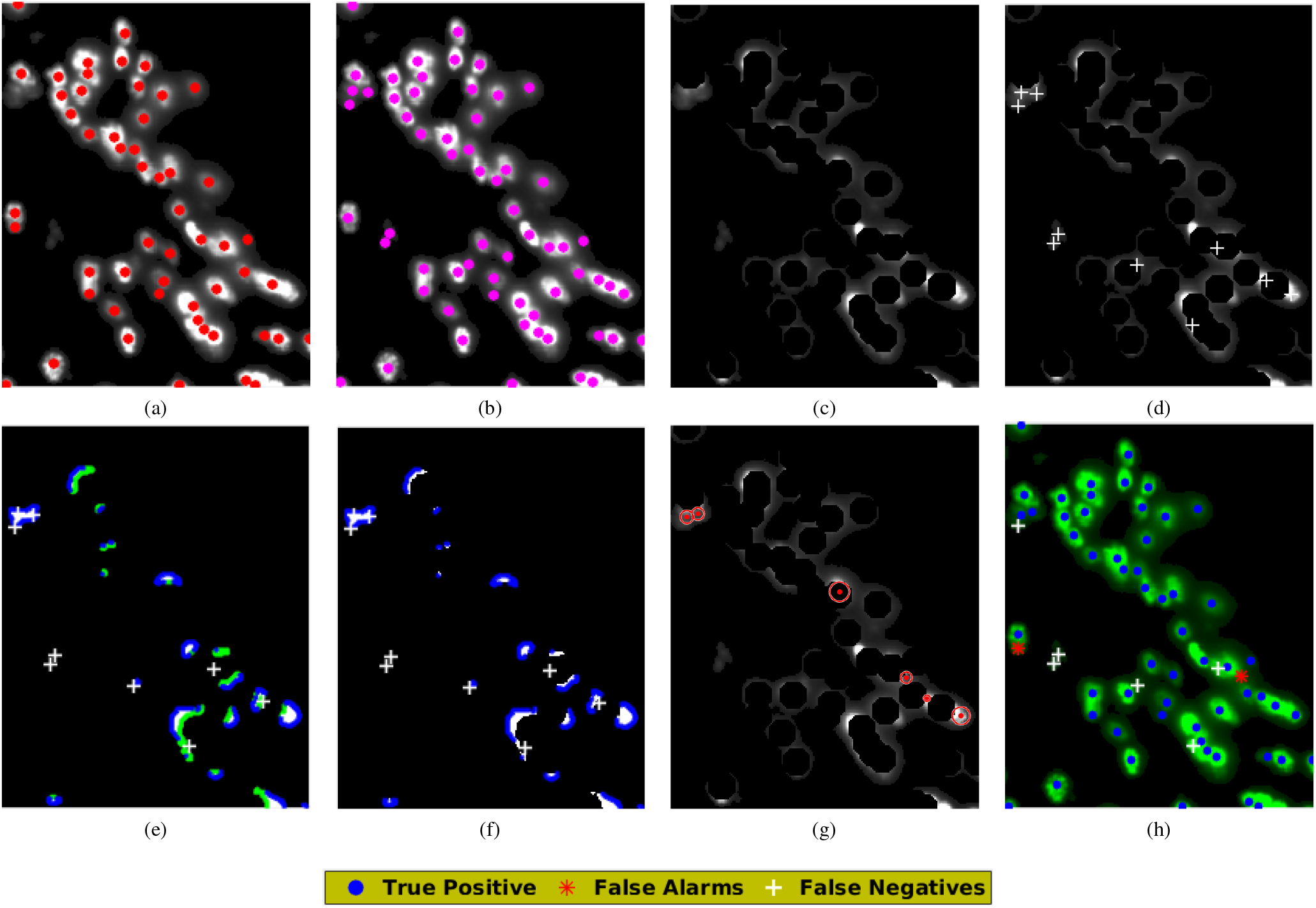
Illustration of the results of the intermediate steps in *Phase-II* sub-processing stage (see algorithm 1 in main manuscript) for the example of strong overlap of GFP nuclei. (a) Centers detected by *Phase-I* of processing on FGR, as given in S.4(g); which serves as the input of Phase II (b) Manually annotated Ground Truth superimposed on the FGR (see Figure 5(a) of manuscript); (c) RFM (Residual foreground map) obtained after the first iteration, from (a); (d) Cells undetected by the *Phase-I* stage of processing, shown superimposed on RFM marked using white crosses; (e) IBC in green and OBC in blue on edges of RFM in (c); (f) filtered (prominent) convex OBCs with undetected set of cells marked in white; and (g) cell centers detected by fitting circles using Hough Transform on convex OBCs in (f); (h) Detected cell centers after *Phase-II*, compared with the ground truth, showing true positives, false alarms, and false negatives; Figure best viewed in color.

**True Positives:** *denote the cell-centers that are detected correctly by our algorithm and are present in the ground truth.*
**False Alarms:** *denote the cell-centers that were missed by our algorithmbut were present in the ground truth.*
**False Negatives:** *denote the cell-centers that were detected by our algorithm but were not present in the ground truth.*

## S.4. Details of Training the supervised ML and DL models used for cell detection, as competing methods

### A. Support Vector Machines (SVM)

For training the shallow supervised methods of classification, we have used annotated cell centers from the ground truth. Using this information, template windows of size 25 × 25 pixels were created over the marked cell bodies. Each window that contains an entire cell body was used as the positive samples for training. We also used other windows from the remaining parts of the brain that contain only partial or no marked cell bodies within the template window which were used as negative samples. Care was taken so as to avoid those windows whose centers are nearby to the labeled cell centers. Examples of positive and negative samples used in the SVM are shown in figure S.6(a) and S.6(b) respectively. The training samples were randomly selected from 3 classified regions as: sparsely distributed, minimal and strong overlap of cells in different parts of brain regions based on density of occurrence of the GFP tagged nuclei. HOG features were then extracted from the samples which were used for training the SVM model. SVM [16] with Gaussian kernel was used for training.

**Fig. S.6:**
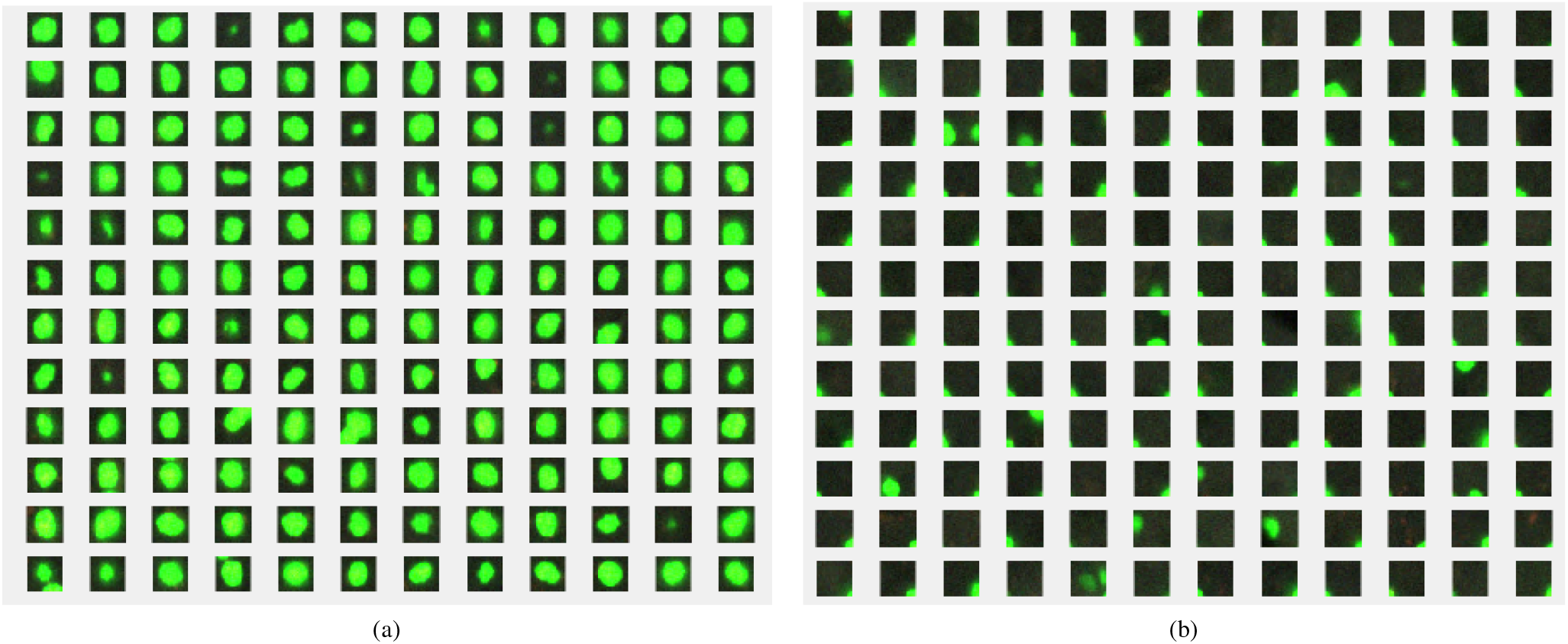
(a) shows the samples consisting of template windows with complete cell nuclei (positive samples) and (b) shows examples with no complete cell nuclei (negative samples) that were used as the the training sets for the SVM [16].

### B. ADABOOST (with HAAR features)

For ADABOOST, we have used features that were extracted from the the mDT map and were used to train the the model. Figure S.7 shows examples of negative and positive samples used for training the ADABOOST model. It is important to note the higher fluorescent irradiance at the center of cells in samples of figure S.7, compared to that in figure S.6. Features, such as, Histogram of Oriented Gradients (HOG) [18] and Multi-Texton Histogram (MTH) [31] are extracted from mDT map (equation 1 in manuscript), for each of the negative and positive mDT samples.

**Fig. S.7:**
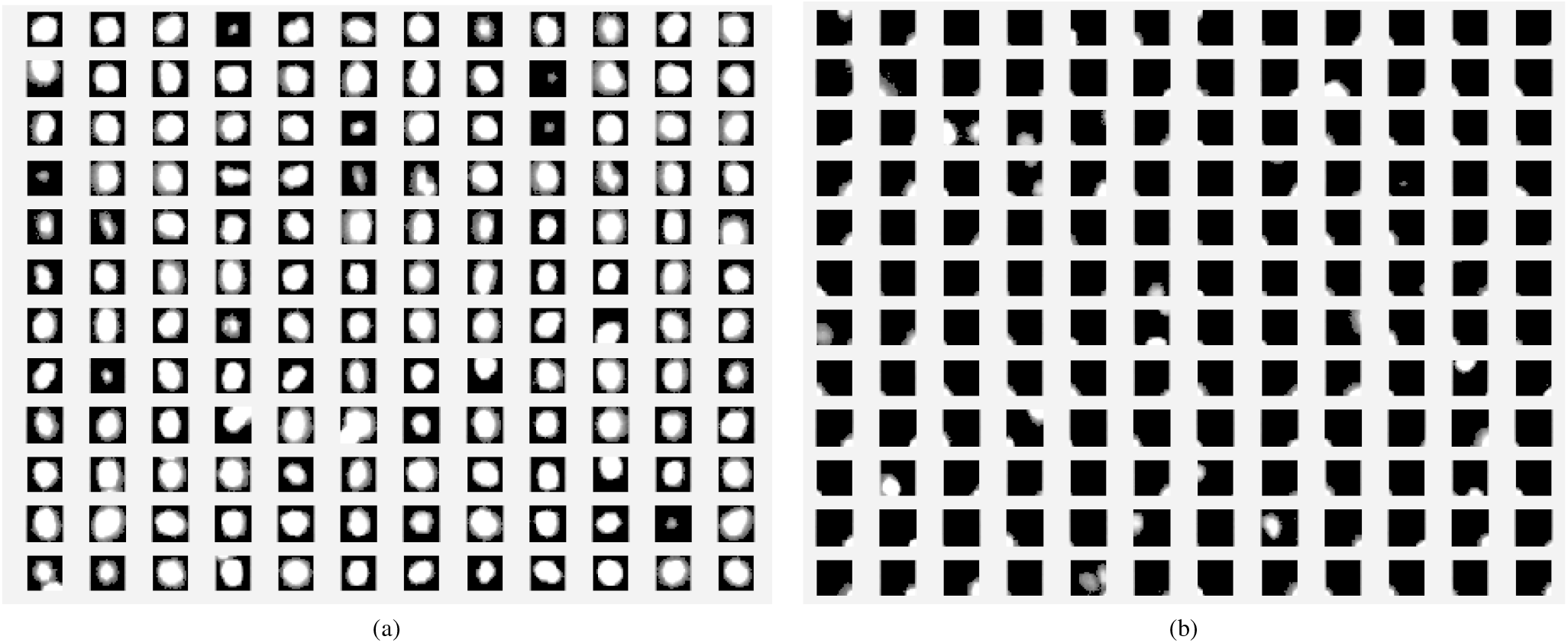
Template samples of mDT obtained from those in figure S.6. These are again used for extracting features, required for training the ADABOOST (with HAAR feature) classifier; (a) *+ve* samples, (b) *-ve* samples.

A total of 25*K* positive samples and 50*K* negative samples were used for training. For training the ADABOOST [57] classifier, Haar features are extracted from the samples. The SVM and ADABOOST implementations of MATLAB are taken off-the-shelf. Positive classification is considered as an area of a detected cell and centers of positive windows were considered as detected cell centers.

### C. CNN models

Two methods of supervised CNN models, Faster R-CNN and SegNet, were also used for experimentation. Faster R-CNN [50] performs a bounding box regression on the objects in an image for detection. We have used equal number of samples for training and testing the Faster R-CNN model. The training stage takes the 224 × 224 pixel images and the coordinates of the bounding boxes around each cell present in the image. A total of 1,47,800 such images are generated each having 2 or more cells in them. We then combine samples from sparse, minimal and strong overlap regions for training the model. Traditionally, Faster-RCNN has been used in literature to detect objects, even when the size varies for each object within an image. Since we have only one object of interest i.e. GFP tagged nuclei in our data, we provide a bounding box of size 25 × 25 pixels around each cell, derived empirically. Faster-RCNN [50] uses the coordinates of the positive samples as the anchor boxes. The negative samples are generated implicitly within the network, i.e. one does not have to provide negative samples to the code as input. Faster-RCNN takes care of the scale and translation invariance. A large number of such anchor boxes are given as input to the Faster R-CNN during the training process and as a result any variability of the cell configuration is taken care during the training. Post training, Faster R-CNN takes an image as input and detects the bounding box around the cell bodies as output. Centroids of the bounding boxes obtained as Faster R-CNN output were considered as detected cell centers. A Tensorflow based implementation of Faster R-CNN [50] is taken of-the-shelf for our experimentation. Samples used as training the Faster R-CNN model is shown in figure S.8. The magenta boxes show the anchor boxes that are fed into the Faster R-CNN network during the training. These anchors are drawn based on the manually annotated data of cell-centers. These square boxes are positioned at these cell-centers with a side length of 11 pixels. Faster-RCNN [24], [50] was trained using a Tensorflow based implementation for 70K epochs. We have used ~ 10^6^ positive training samples for Faster R-CNN (compared to [50], which reports using 105 samples). The anchor boxes are equivalent to those in PASCAL-VOC challenge; where the objects are simple and different. In our case, each window contains an entire cell body and were labeled as positive samples. The Faster R-CNN techniques used for COCO (image classification) do not use more than 10K samples per class. Since the problem at hand is a two-class problem and the training set size was ≃ 10^6^, we aimed to find a trade-off between the training time and the number of samples. We also found that increasing the samples 3-fold did not significantly increase the performance of the system. Our Faster R-CNN parameter count is 2.4 × 10^6^ [50]. Our model was trained for 7 × 10^6^ epochs using the codes provided by the authors [50].

For the problem at hand, recent literature in the domain of Deep Learning does not have an unsupervised framework for CNNs, to the best of our knowledge. Hence, we have resorted to comparing our results with the supervised CNN architectures.

**Fig. S.8:**
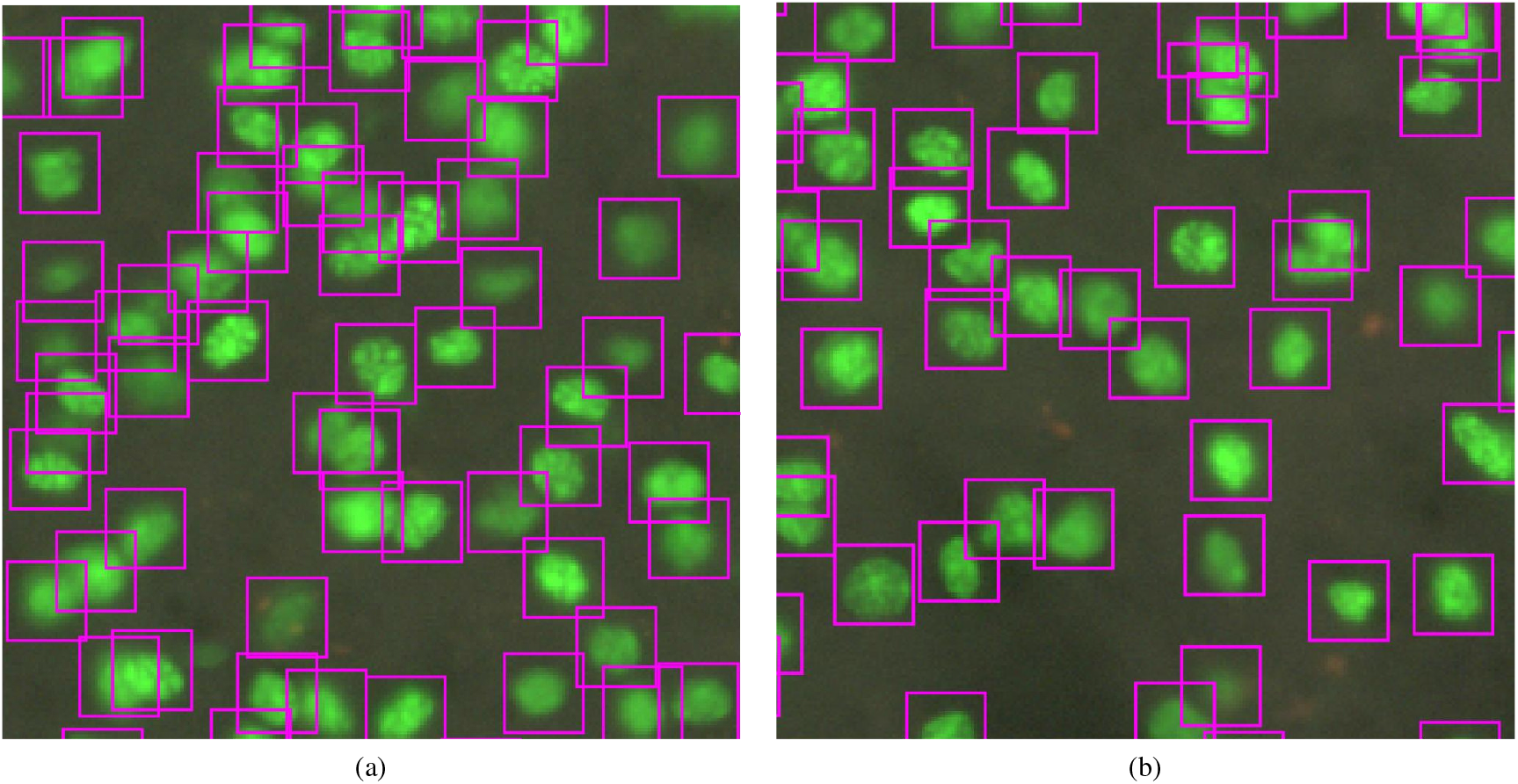
Two sample sections of the brain image used for training the Faster R-CNN [50]. Bounding boxes are positioned using human-annotated groundtruth data. No negative samples needed.

The SegNet [6] model has been trained as per the code provided by the authors. It’s a supervised (CNN) method used previously for detecting cells in blood. Training samples (positive ones only needed) for SegNet were generated by delineating circular patches around manually annotated cell centers. In this case, we used a circular patch with the radius of 6 pixels around the manually annotated cell-centers, as inputs for training. The trained SegNet map produces outputs as circular patches (sometimes overlapping with each other) on the test image, whose centers are detected as cell-centers. The SegNet [6] is a torch based implementation that is used previously for segmentation in computer vision. The SegNet model is a Variational auto-encoder based model, which inputs RGB image at one end and its binary segmented map on the other end. Based on the annotated cell-centers, circular patches with radius (*r* = 6 pixels) are inpainted to create virtual segmented maps of the original image, for training the SegNet model.

**Fig. S.9:**
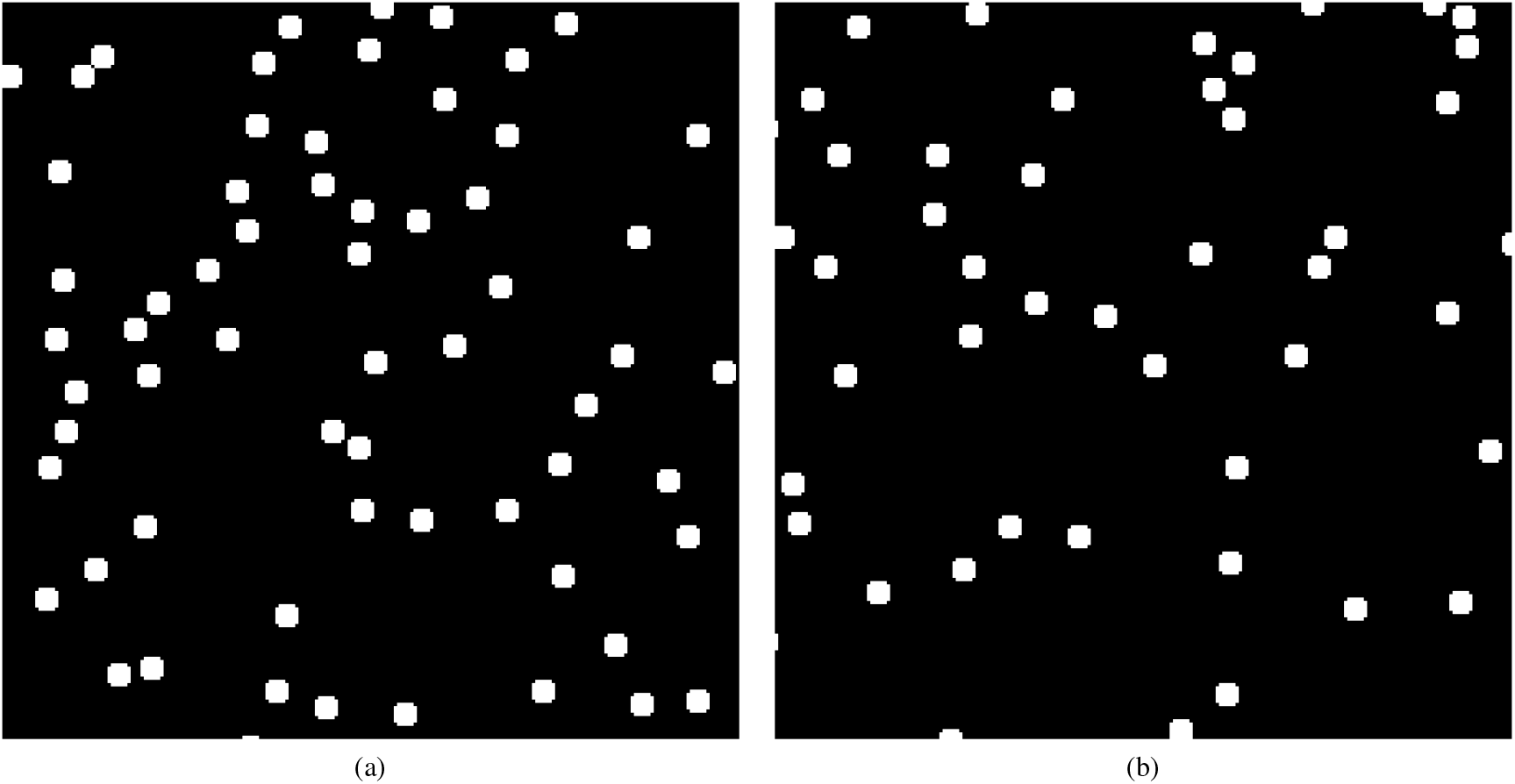
Two sample sections of the brain image used for training the SegNet [6]. Circular patches are drawn using human-annotated ground truth data. No negative samples needed.

**TABLE S.2:**
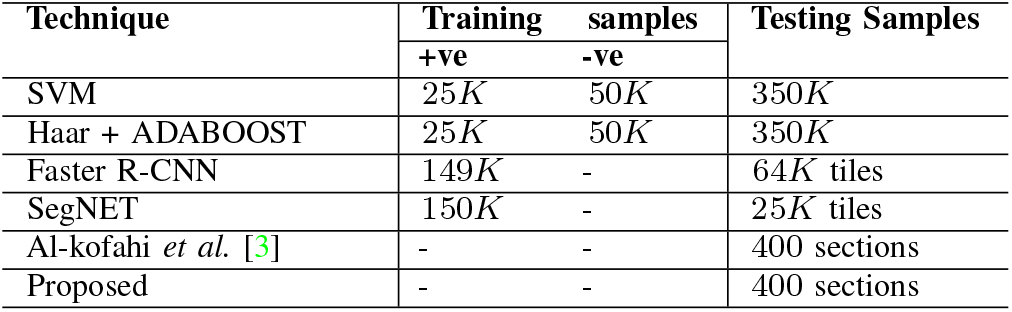
Number of samples used for training and testing (each tile is a cropped section of brain scan of size 512 × 512 pixels).

## S.5. Use of NMS to generate final cell locations using supervised models

Implementations of Faster R-CNN and ADABOOST (using HAAR features) classifiers produce multiple bounding boxes, while SVM produces multiple candidate locations as target outputs during the testing phase. These positive responses (output candidates) obtained from the trained machines, typically appear as clusters (of strong overlapping boxes in cases of Faster R-CNN and ADABOOST). SVM implementation, on the other hand, produces clusters of positive responses with varying confidence levels. The same is true for the other two methods, where each output appears with a confidence score along with a scale of the bounding box.

Most implementations of Faster R-CNN, ADABOOST and SVM use a method of post-processing (based on local averaging) on these nearby candidates, to obtain the final target position. These implementations did not produce satisfactory results and have thus been modified by us. The primary reason for this failure is that, in general their design does not deal with our scenario of minimal and strong overlapping presence of multiple identical objects (e.g. thin crowd, swarm of bees, ant colony etc.) as targets [51].

To obtain a better result for finalization of the target (cell) locations, we have used a Non-maximal Suppression (NMS) [54], [9] strategy. The solution demands the detection of a maximal (local) response from a 2-dimensional clutter of close-by responses with varying confidence scores. We have used a minor modification of the post-processing from the implementation of [22], as a greedy method of NMS. The modification involves: Within a large neighborhood, *W*, instead of averaging the overlapping window responses, we consider a rank-ordered list of candidates sorted by the confidence scores. While doing so we eliminate outliers of extremely small and large scales (range learned using responses from GT data). Lastly, choose the topmost confidence scores as winner (max) within the local window (*W*), and update the list to apply this last step of NMS iteratively till there are no further significant peaks of confidence score left as a candidate response. This method works reasonably well for our specific scenario of minimally and strongly overlapping objects (brain cells). The qualitative (visual) results of the supervised processes: SVM, HAAR+ADABOOST and Faster R-CNN, along with their evaluation studies given in tables 1-3, are all hence reported using the outputs of this NMS-based process.

The outputs obtained from the software implementation of Faster R-CNN [24], [50] are available as bounding boxes on the test images. The centers are calculated for each of these square boxes to get the cell-centers. A similar strategy is used for the output of SegNet, which produces the segmented maps as circular blobs, where the center for each blob is taken as the detected cell-center. The next section gives qualitative comparison of the results of all unsupervised and supervised methods using two strong overlap and one minimal overlap regions.

## S.6. Additional Results and Extended Discussions

This section exhibits a few additional results of different (competing) methods used to get the cell-centers. The following figures show the output of competing methods (see table S.2) on two overlap and one minimal overlap regions of cells in the mouse brain scan. Results of the proposed method appear in main manuscript (figures 9 and 10).

**Fig. S.10:**
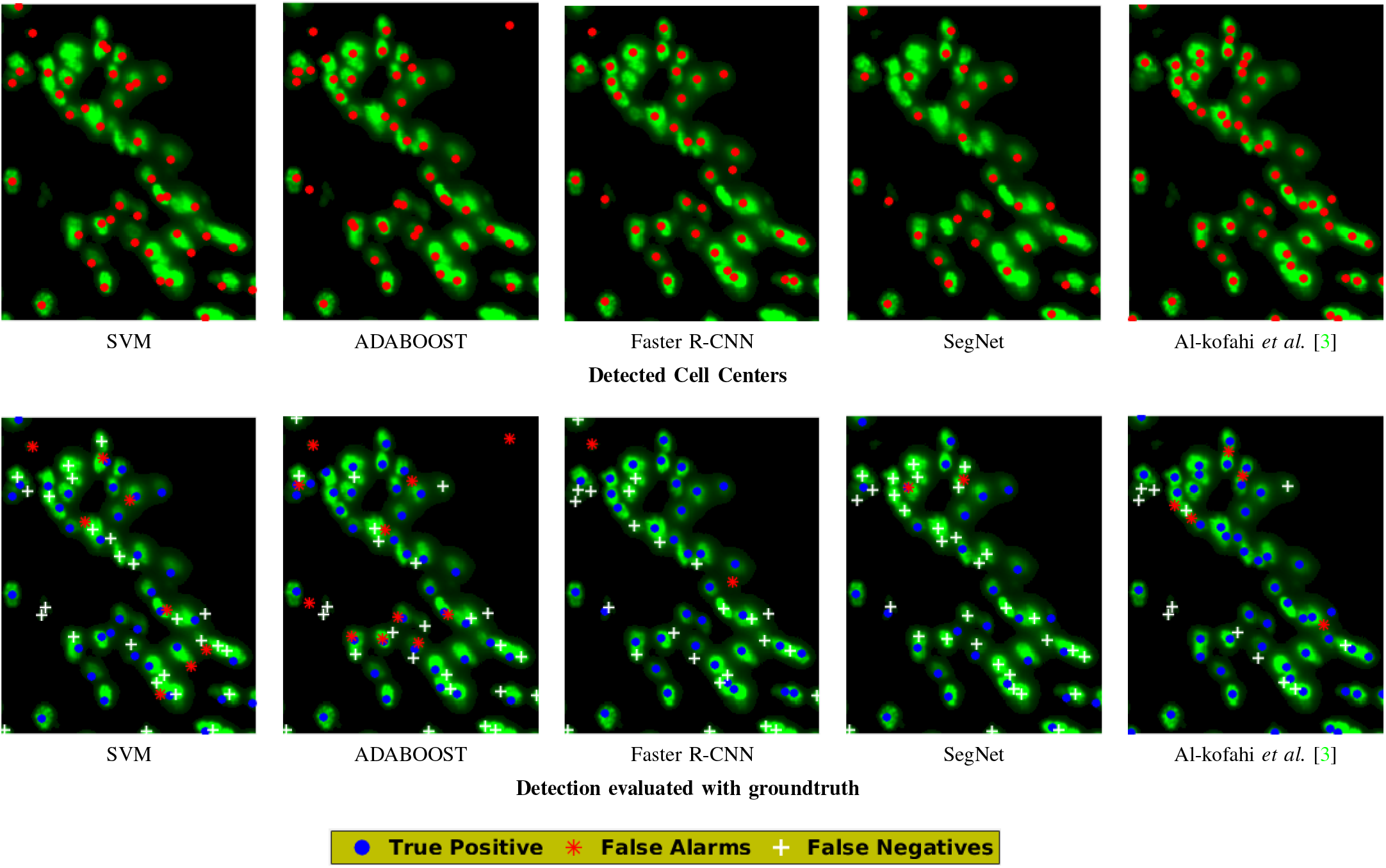
Strong Overlap Region: Visual illustrations for qualitative comparative study of performance of different competing methods, for the image given in Figure 3(d) in main manuscript (best viewed in color).

Section II-A in main manuscript gives the basis and justifies the design of our proposed unsupervised pipeline. Other alternative strategies (for *Phase-II*) which we had pondered and partly explored for the solution to the problem of cell detection are: (i) 2-D bin packing problem; (ii) Minimize an optimization function for placement of cells, with suitable scores and cost criteria; and (iii) image saliency based on superpixels [2]. A few trials did not give us encouraging results (*e.g.*, trapped in a false minima while minimizing), which were worse than even those produced by the simplest Gaussian convolution (see table 1, section VI-A in manuscript) operation. We are certain that this dataset will throw theoretical and analytical challenges in overlapping fields of image processing (and CV), computational geometry and variational optimization.

One may have an impression that the inter-neuron somata are easier to detect than pyramidal cell somata. This is not necessarily true: the difficulties here have to do with the degree of overlap of objects in the image, which depends on cell densities as well as microscopic methodology. The previous results, published in [28], only reach 90% performance level (in contrast with our performance levels of > 97%), something that is easily seen on visual examination of brain images from [28] (not shown here). The proposed algorithm is completely automated and unsupervised in its application.

**Fig. S.11:**
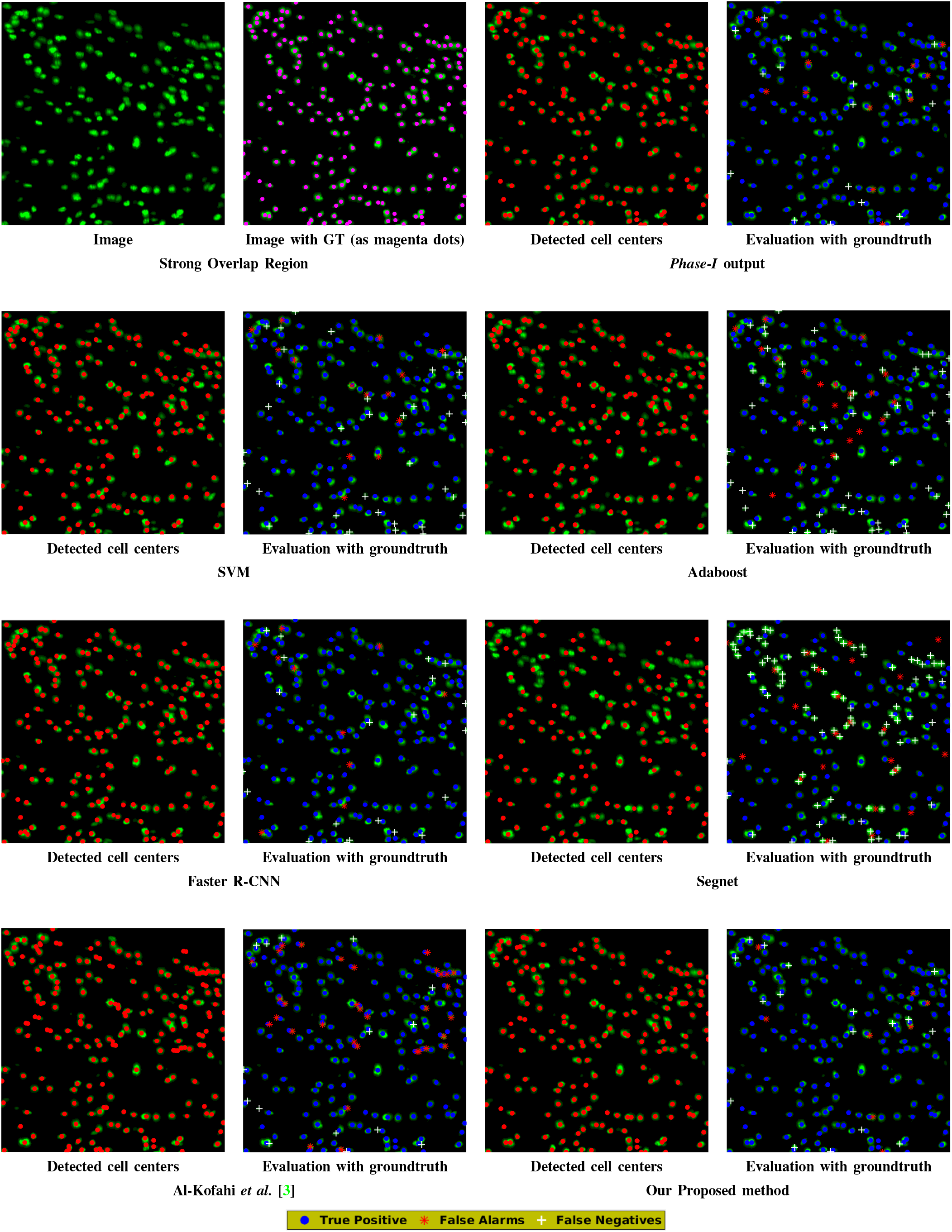
Strong Overlap Region: Visual illustration for qualitative comparative study of performance of different competing methods, for the image given in Figure 10(c) in main manuscript (best viewed in color).

**Fig. S.12:**
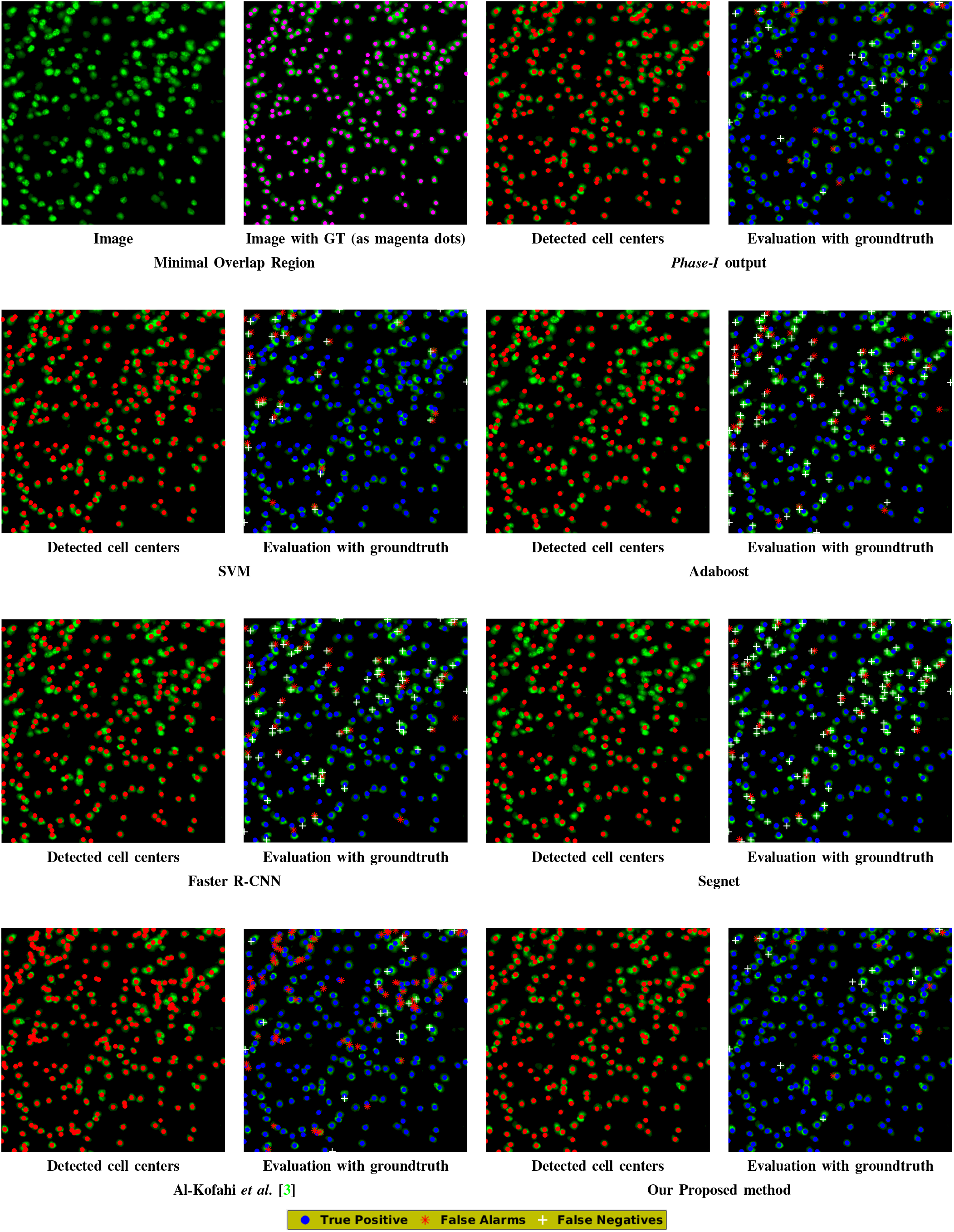
Minimal Overlap Region: Visual illustration for qualitative comparative study of performance of different competing methods, for the image given in Figure 10(a) in main manuscript (best viewed in color).

## S.7. Samples of GUI interface in the tool used for manual annotation

Figures S.13 and S.14 show the screenshots of the GUI for the manual annotation tool, used to obtain ground-truth.

**Fig. S.13:**
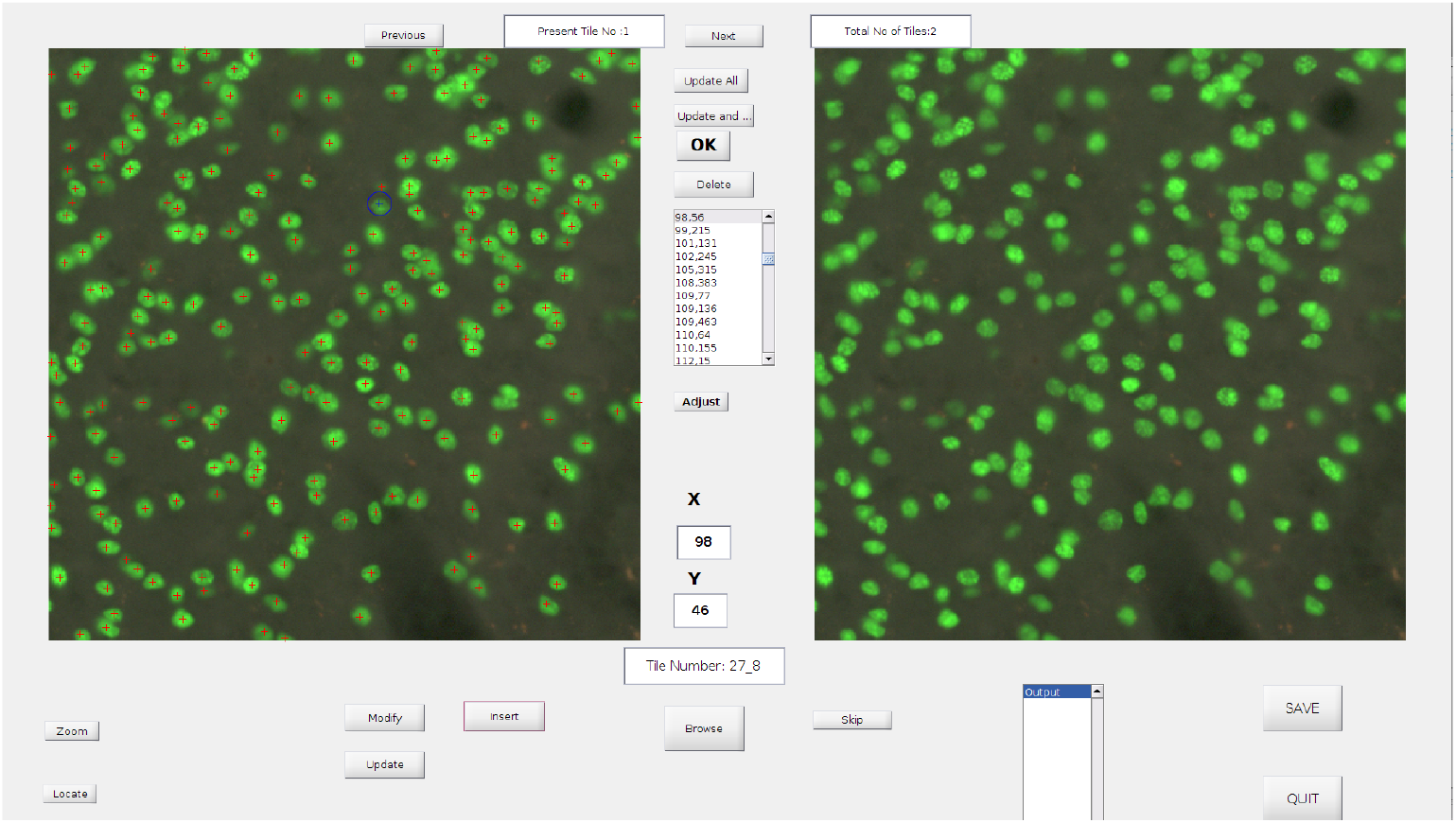
Illustration of the manual annotation tool over a minimal overlap region of the brain scan.

**Fig. S.14:**
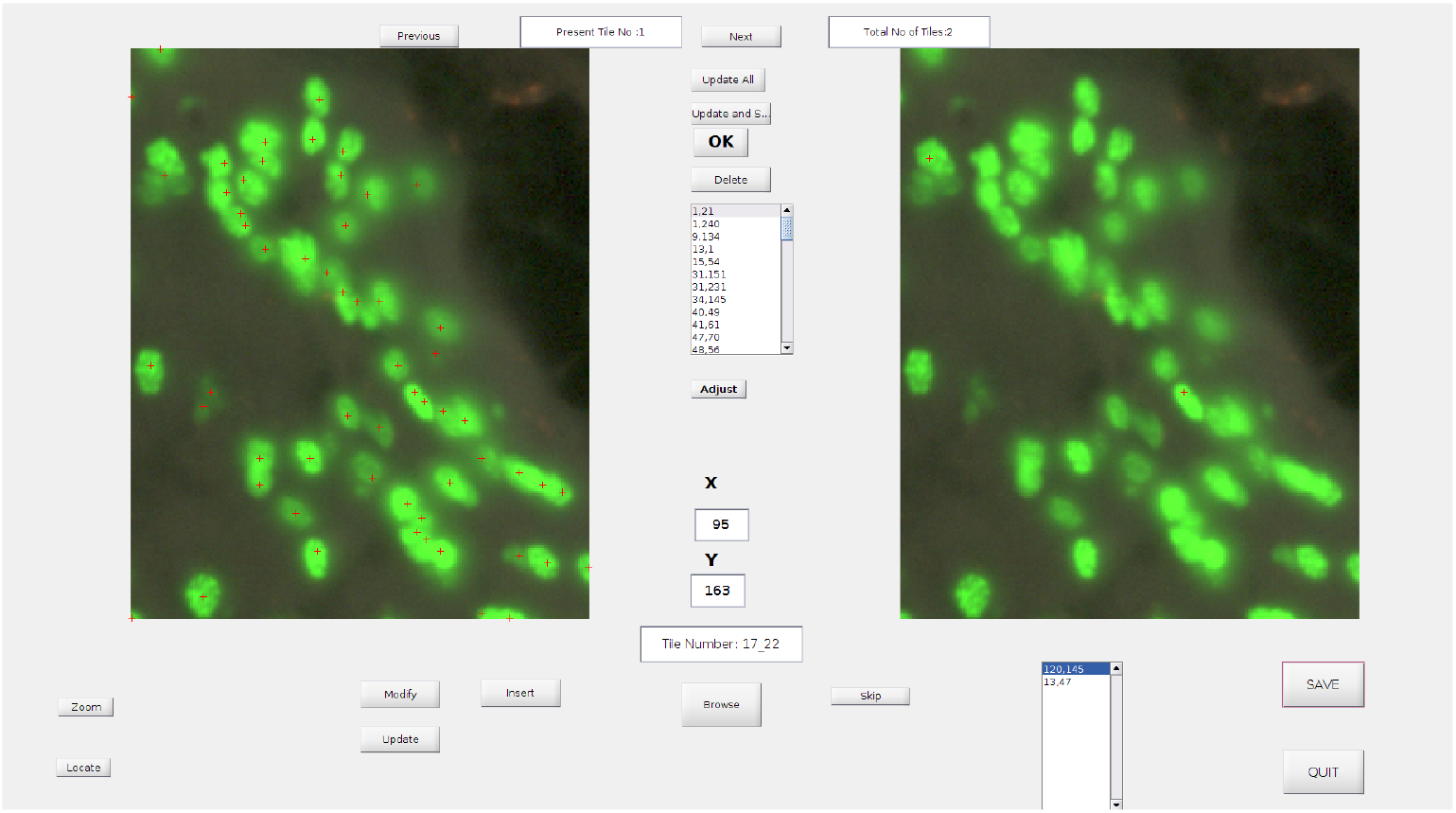
Illustration of the manual annotation tool over a region of cell overlap in the brain scan.

